# Rif1 prolongs the embryonic S phase at the *Drosophila* mid-blastula transition

**DOI:** 10.1101/231357

**Authors:** Charles A Seller, Patrick H O’Farrell

## Abstract

In preparation for dramatic morphogenetic events of gastrulation, rapid embryonic cell cycles slow at the Mid-Blastula Transition, MBT. In *Drosophila melanogaster* embryos, downregulation of Cdk1 activity initiates this slowing by delaying replication of satellite sequences and extending S phase. We found that Cdk1 inhibited the chromatin association of Rif1, a candidate repressor of replication. Furthermore, Rif1 bound selectively to satellite sequences following Cdk1 downregulation at the MBT. In the next S phase, Rif1 dissociated from different satellites in an orderly schedule that anticipated their replication. Rif1 lacking potential phosphorylation sites failed to dissociate and dominantly prevented completion of replication. Loss of Rif1 in mutant embryos shortened the post-MBT S phase, and rescued embryonic cell cycles disrupted by depletion of the S phase-promoting kinase, Cdc7. Thus, *Drosophila* Rif1 mediates the MBT extension of S phase and functionally interacts with S phase promoting kinases to introduce a replication-timing program.

## INTRODUCTION

Eukaryotic DNA replication begins at many locations throughout the genome known as origins. Different origins initiate at different times during S phase on a schedule governed by an elusive replication timing program. The time it takes to duplicate the genome, the length of S phase, is set by the time when the last sequence completes replication. For over 50 years, the field has appreciated that late-replicating sequences are found in the compacted portion of the genome known as heterochromatin (Lima-De-Faria 1959; Lima-De-Faria and Jaworska 1968; Taylor 1960). Late replication is presented as a general property of heterochromatin, but how this property arises is unknown (Lima-De-Faria and Jaworska 1968). Additionally, embryonic development of many animals features dramatic modifications of replication timing (Graham and Morgan 1966; Snow 1977; Stepinska and Olxanska 1983; Hiratani *et al.* 2008). In *Drosophila,* the length of S phase changes by over 50 fold during development (Blumenthal *et al.* 1974; Neufeld *et al.* 1998; Milan *et al.* 1996). How such alterations are made is unclear.

As worked out in yeast, the process of origin initiation involves a sequence of conserved biochemical steps that convert an origin into a bidirectional replication fork (Tanaka and Araki 2013; Yeeles *et al.* 2015). Origins are first licensed for replication through the loading onto dsDNA of two hexameric MCM2-7 helicase complexes. The resulting head to head double hexamer is known as a pre-Replicative Complex (pre-RC). Activation of the pre-RC requires the coordinated assembly of a multiprotein complex called a replisome. Activation, also referred to as firing, is led by the action of two conserved cell cycle kinases: a CDK (cyclin-dependent kinase) and a DDK (Dbf4-dependent kinase) (Labib 2010). CDK and DDK are recruited to pre-RCs where they phosphorylate essential substrates to initiate the transformation of the pre-RC into functional replication forks (Tanaka *et al.* 2006; Zegerman and Diffley 2006; Sheu and Stillman 2010). Temporal aspects of the cell cycle, and local chromatin structure are thought to influence this step to govern the replication-timing program (Rhind and Gilbert 2013). Multiple studies have identified a number of pathways contributing to replication timing including local factors, such as histone modifications (Vogelauer *et al.* 2002; Schübeler *et al.* 2002; Aggarwal and Calvi 2004) and chromatin binding proteins (Makunin *et al.* 2002; Knott *et al.* 2012), and global factors, such as the competition among origins for limited replication components (Tanaka *et al.* 2011; Mantiero *et al.* 2011). How different factors contribute to the length of S phase and the phenomenon of late replication has been difficult to resolve likely due to their redundant effects in differentiated cells.

In light of these issues, the embryo provides a unique context in which to study the control of S phase duration because development profoundly changes it. Historically, the study of cell cycle timing has focused on the G1/S and G2/M transitions, but the early embryonic cell cycles lack gap phases and it is the duration of S phase that changes (McCleland *et al.* 2009; Shermoen *et al.* 2010). In the *Drosophila melanogaster* embryo, these specialized cell cycles are maternally programmed, synchronous, and occur in the absence of cell membranes. During the first 9 nuclear cycles S phase completes in as little as 3.4 min (Blumenthal *et al.* 1974), and nuclei remain deep within the embryo. Beginning in cycle 10 the nuclei reach the surface of the embryo forming the blastoderm. During cycles 11-13 the length of S phase gradually increases to 13 min. At the next cell cycle, the 14^th^, the embryo begins a conserved developmental period known as the Mid-Blastula Transition (MBT) when numerous changes occur: interphase, which extends dramatically, has an S phase 14 that lasts 70 minutes and it is followed by the first G2 (summarized in Figure 1A) (Farrell and O’Farrell 2014; Yuan *et al.* 2016). After G2 of cycle 14, cells enter mitosis in a patterned program controlled by zygotic transcription (Edgar and O’Farrell 1989 and 1990). Our focus is on the changes that occur to the DNA replication program at the MBT, and on how these changes are introduced.

**Figure 1.**
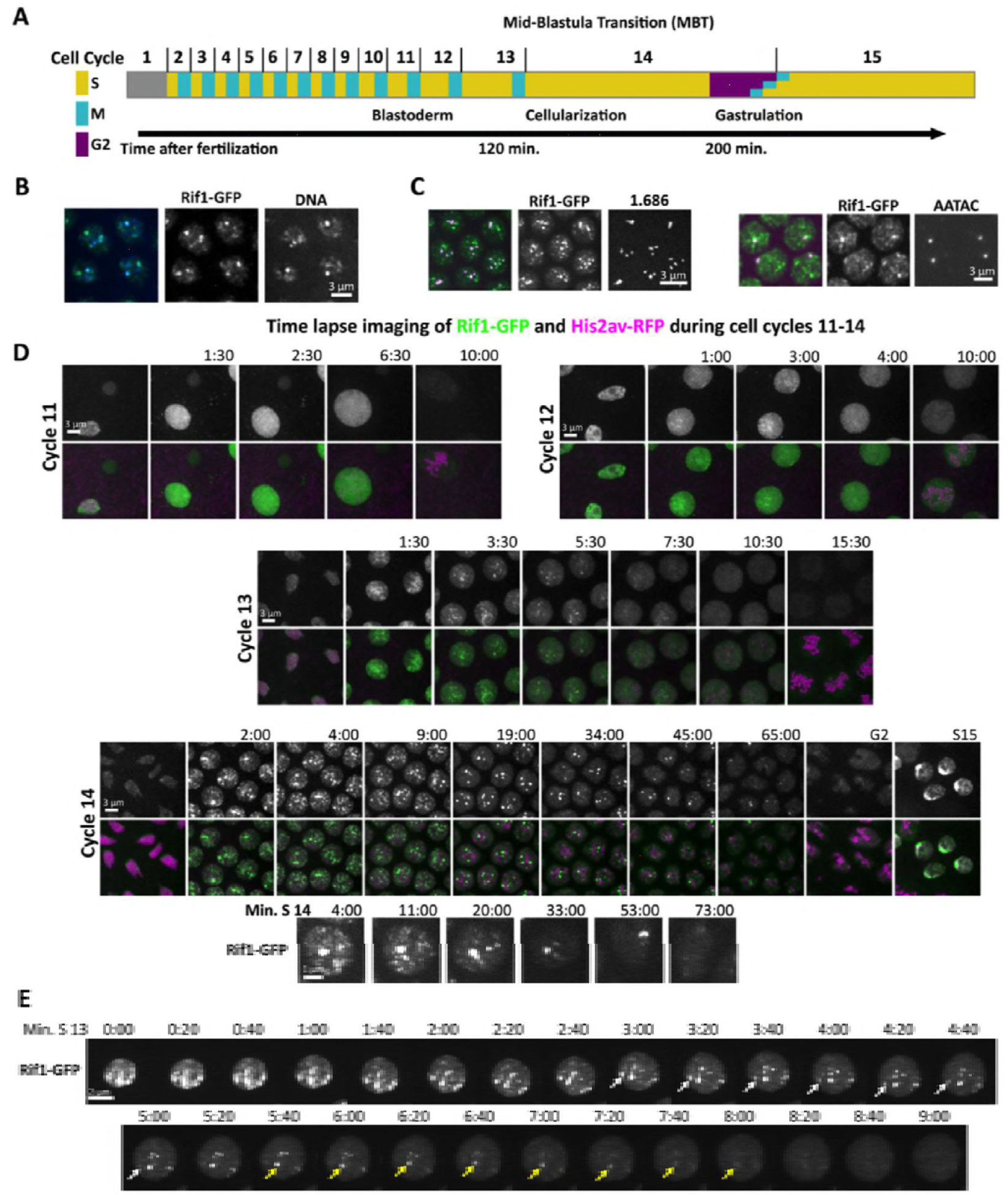
Rif1 forms foci on heterochromatin as the embryonic S phase lengthens. (A) Diagram highlighting the key changes to the cell cycle and to embryonic morphology during early development. The initial prolongation of the cell cycle is due to the increase in the length of S phase. This occurs gradually during cycles 11-13, and then the onset of a late replication program in S phase 14 extends interphase considerably. The first G2 is introduced in cycle 14, after which cells enter mitosis 14 asynchronously, according to a developmental pattern. **(B)** Nuclei from an S phase 14 Rif1-GFP embryo stained for GFP (green in the merge) and DNA (DAPI: blue in the merge). Rif1 co-localizes with regions of the interphase nucleus that stain intensely with DAPI. **(C)** Specific satellites (magenta in the merge), either 1.686 detected live by TALE-light or AATAC detected by DNA FISH co-localized with Rif1-GFP (green in merge) during early cycle 14. **(D)** Frames from live imaging (Movie 1) showing Rif1-GFP and H2aV-RFP in developing embryos during cell cycles 11-14. Note the initial presence and increasing persistence of nuclear Rif1 foci as interphase lengthens. Below, magnified images from time-lapse records of Rif1-GFP during S phase 14. Individual nuclear foci of Rif1 disappeared at different times across S phase. **(E)** Frames from time lapse showing Rif1-GFP in a single nucleus at 20 sec intervals during S phase 13 (from Movie 2). Arrows indicate two specific foci at the time that their fluorescence declined.

Prior work has shown that the prolongation of S phase at the MBT is caused by reduced synchrony in replication timing (Shermoen *et al.* 2010). During the pre-blastoderm cycles all regions of the genome begin and end replication together resulting in a short S phase. Even the 30% of the genome composed of blocks of repetitive DNA known as the satellite sequences, which are considered to be constitutively heterochromatic, lack the marks of heterochromatin and replicate early. During S phases 11-13, the satellite sequences experience progressively modest delays in replication timing. Then, these satellites experience major delays in S phase 14 in which a succession of different satellites begin and complete replication on a protracted schedule. Thus, onset of late replication causes the pronounced increase in the S phase duration. Following their late replication in cycle 14, the satellite sequences become heterochromatic and late replication will be a characteristic feature of heterochromatic satellites for the rest of development.

Uncovering how late replication develops will improve our understanding of what has been a widespread, but mysterious feature of DNA replication. We know that the onset of late replication in the embryo follows a time-course that is unrelated to appearance of other features of heterochromatin such as compaction, which is already evident in earlier cell cycles, or HP1 recruitment, which occurs after replication in cycle 14, or H3K9 methylation, which accumulates continuously and slowly over the course of interphase 14 (Shermoen *et al.* 2010; Yuan and O’Farrell 2016). Previous work demonstrated that activity of the Cdk1 kinase is a key determinant of the duration of S phase in the early embryo (Farrell *et al.* 2012). Persistent S phase Cdk1 activity during the shorter pre-MBT cycles drives heterochromatin to replicate early, and it is the programmed downregulation of Cdk1 occurring at the MBT that allows the onset of late replication. However, it is unclear how the embryo interprets the activity of Cdk1 to produce this dramatic prolongation of S phase.

Here we focus on the conserved protein Rif1 because recent work in yeast and vertebrate cells indicates that Rif1 can regulate the timing of origin firing (Hayano *et al.* 2012; Cornacchia *et al.* 2012; Yamazaki *et al.* 2012). Rif1 interacts with the protein phosphatase PP1 through conserved sites and recruits PP1 to chromatin where it can play roles in telomere biology, DNA damage responses and control of replication (Mattarocci *et al.* 2016). Several studies in yeast, and more recently in vertebrates, suggest that Rif1 recruited PP1 opposes the action of DDK by dephosphorylating this kinase’s essential substrate, the MCM helicase complex (Mattarocci *et al.* 2014; Davé *et al.* 2014; Hiraga *et al.* 2014; Alver *et al.* 2017). In support of this model, deletion of yeast *rif1* can partially rescue viability to lethal mutants of *cdc7* or *dbf4* (Hayano et al., 2012). How Rif1 is itself inhibited during S phase, thereby allowing late origins to initiate, is unclear, although the involvement of DDK and CDK has been suggested. Additionally, how Rif1 contributes to developmental changes in replication timing is unknown. Our results describe how Rif1 connects the activity of Cdk1 to the length of the embryonic S phase, and provides new insights into the regulation of Rif1 during the cell cycle, and during development.

## RESULTS

### Rif1 binds to compacted foci of satellite sequences as the length of the embryonic S phase increases

Due to the stereotyped nature of the embryonic cell cycles, live imaging of fluorescently tagged proteins can be especially informative. We used CRISPR-Cas9 genome editing to tag the endogenous Rif1 protein with EGFP at its C-terminus (Supplemental Figure 1). Rif1 is maternally provided (Supplemental Figure 1C), and is widely distributed during the first 6 hours of embryogenesis and thereafter shows increasingly tissue-limited expression (Sreesankar *et al.* 2015).

Given ubiquitous presence of Rif1 protein in the early embryo, regulation of its activity ought to underlie any developmental or cell cycle modulations of Rif1 function at this stage. If Rif1 acts to delay the replication of heterochromatin, then we would expect the protein to be recruited to satellite sequences when the replication of these sequences is delayed in cycle 14. The satellite sequences form compacted regions of chromatin that can be visualized as discrete bright foci of DAPI stained DNA (Figure 1B). These compacted foci of chromatin first acquire heterochromatic marks during cycle 14, but, given the absence of G1 in this cycle, a program for their delayed initiation must already be in place at the beginning of interphase 14 (Shermoen *et al.* 2010; Yuan and O’Farrell 2016). In S phase 14, Rif1 was bound to many foci of compacted chromatin, while in the following G2 phase the heterochromatin lacked Rif1 staining (Figure 1). Live imaging of Rif1-EGFP embryos (Movie 1) during early embryogenesis revealed the dynamics of this change. Rif1-EGFP disappeared from individual foci as S phase progressed, and only a dim nuclear background was present by G2 of cycle 14 (Cycle 14, Figure 1D). Tracking individual foci in high frame-rate movies showed that different Rif1 foci disappeared at different times (bottom panels, Figure 1D). Often, only a single focus remained near the end of S phase before disappearing as cells entered G2. This progressive loss of different Rif1 foci was reminiscent of the protracted schedule of late replication occurring in this cycle.

To test the correspondence of Rif1 foci and satellite sequences we marked specific satellite sequences by in situ hybridization or our recently developed TALE-lights technique (Yuan *et al.* 2014). In embryos fixed during early S phase 14, the single Y-chromosome linked focus of AATAC satellite detected by DNA-FISH was co-stained by Rif1 (right panels Figure 1C). In live embryos, a fluorescently tagged TALE-light protein engineered to bind the late-replicating satellite-repeat 1.686 showed that Rif1 also binds the four foci of the this sequence (left panels in Figure 1C).

During cycles 11-13, S phase gradually and progressively gets longer due to incremental delays to satellite replication. In paralleled with these changes, Rif1 showed weak and transient localization to foci in cycle 11 and progressively more intense and longer-lived foci in subsequent cycles (Figure 1D). Disappearance of Rif1 foci within each S phase was not instantaneous: the signal decayed from the outside-in over a few frames of our records, for instance during S phase 13 high frame imaging showed that a focus of Rif1 disappeared over the course of approximately 2 minutes (Movie 2, Figure 1E). These observations show that Rif1 binds to satellite sequences as they become late replicating during embryogenesis, and that Rif1 dissociates from satellite sequences as replication progresses.

### The release of Rif1 from chromatin anticipates the initiation of late replication

The correlation between the dissociation of the Rif1 localized to satellite foci and the onset of late replication motivated a closer examination of Rif1 dynamics during S phase. We have previously validated and utilized fluorescently tagged versions of the replication protein PCNA as real time probes for the progress of S phase (McCleland *et al.* 2009). PCNA travels with the replicating DNA polymerase, and its recruitment to different regions of the genome marks their replication. Using a transgenic line expressing an mCherry labeled version of PCNA from its endogenous promoter (Supplemental Figure 2), we were able to compare active DNA replication with the localization of Rif1. During the first 10 minutes of S phase, the PCNA signal was intense, and distributed throughout most of the nucleus. Following early replication, the generalized signal fades progressively and the PCNA signal is predominantly limited to bright apical foci, which mark the late replicating satellite sequences. Each late replicating focus appears, persists, and declines in a stereotyped schedule, with the number of active foci gradually declining during more than an hour of interphase of cycle 14 (Figure 2A, Movie 3). Throughout this program, Rif1 did not overlap with the PCNA signal, indicating that once a region had initiated DNA replication, it no longer bound obvious levels of Rif1. As S phase progressed, different Rif1 foci disappeared at different times and were replaced within one minute by PCNA (Movie 3). The last sequence to recruit PCNA was marked by Rif1 throughout S phase 14 until just before it recruited PCNA (Figure 2A). Similar analyses in cycles 12 and 13 similarly revealed separation of Rif1 staining and replication. The observed timing suggested that replication of Rif1 staining regions awaits the dissociation of Rif1.

**Figure 2.**
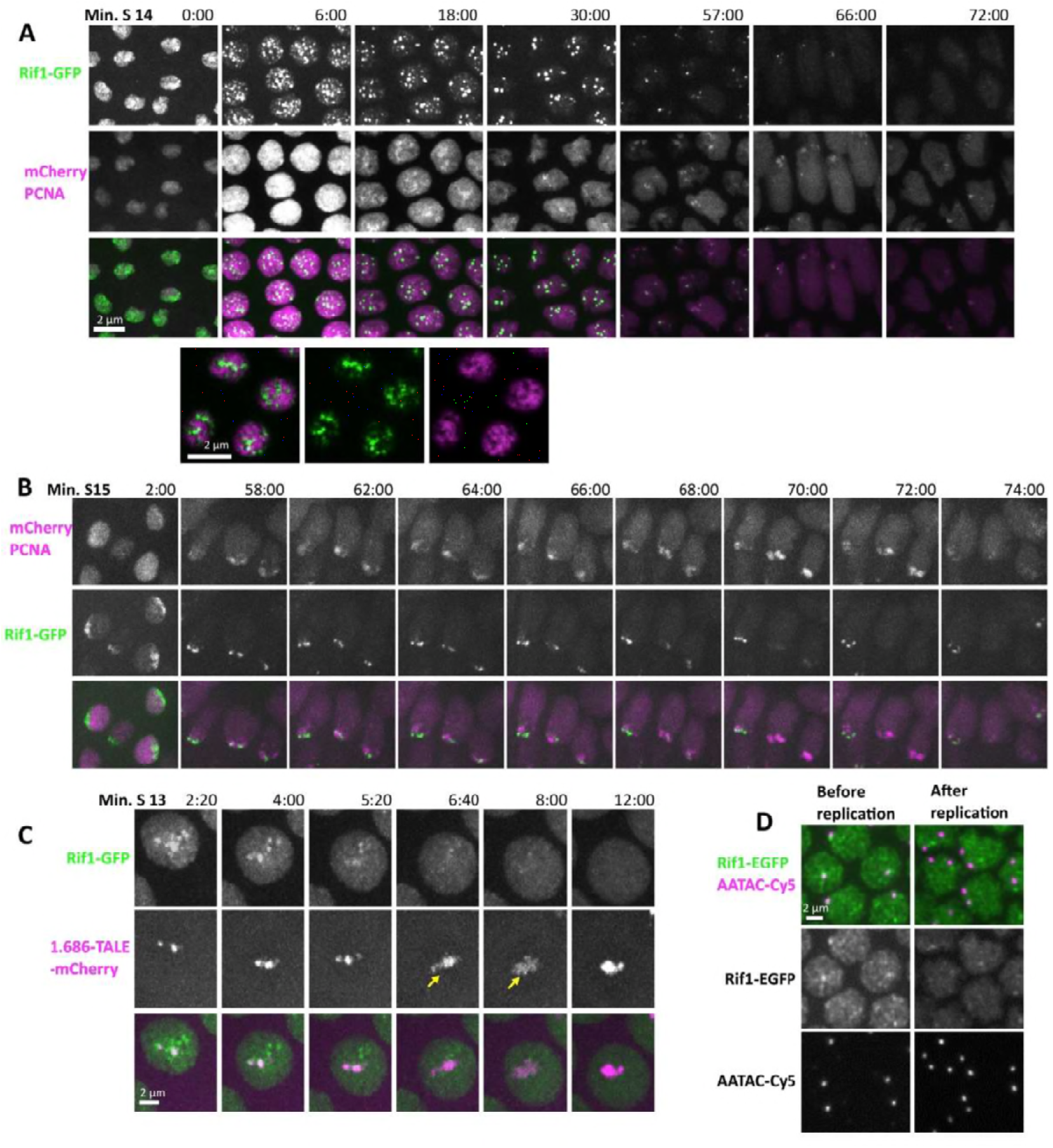
Rif1 dissociates from chromatin before the underlying sequences replicate. **(A)** Stills from time-lapse microscopy of Rif1-GFP and mCherry-PCNA during S phase 14 (Movie 3). Though both form foci on satellite sequences, the signals do not overlap. Rif1 marks numerous satellites early and the number of foci progressively declines. After early widespread staining, PCNA, a marker of active replication, transiently marks a progression of different satellites. At different positions, the period of PCNA staining seems to follow loss of Rif1. Below, a single z plane from 10 min into S phase 14 that more clearly shows that the signal from Rif1 does not overlap with that from PCNA. **(B)** Stills from timelapse microscopy of Rif1-GFP and mCherry-PCNA during S phase 15 (Movie 4). During this S phase the late replicating heterochromatin is clustered into an obvious chromocenter. As in S phase 14, Rif1 marks late replicating chromatin, and Rif1 dissociates from these sequences before the acquisition of PCNA. **(C)** Live imaging of Rif1 and the satellite 1.686 using an mCherry tagged TALE protein (Movie 5). Yellow arrow indicates the decompaction and replication of 1.686. After its replication, the 1.686 repeat re-compacts and appears brighter. Rif1 dissociates from this sequence prior to its replication. **(D)** Nuclei in S phase 14 stained for Rif1 and for the Y-chromosomal satellite repeat AATAC. Rif1 is bound to AATAC during early S14 before the AATAC repeat replicates. During late S14 the duplicated AATAC sequence lacks Rif1 stain.

The S phase of cell cycle 15 provides an alternative context to observe the relationship between Rif1 and replication. By cycle 15, the embryo has completed the MBT, the cell cycle has lost synchrony and morphogenetic movements have begun. The timing of mitosis 14 and entry into cycle 15 differs with position in the embryo, but follows a stereotyped schedule that is zygotically controlled. Cycle 15 still lacks a G1, so nuclei enter S phase immediately following mitosis 14, but these cycle 15 nuclei exhibit more mature features. The satellite sequences enter cycle 15 after introduction of heterochromatic marks such as H3-K9me and localized HP1 during cycle 14. Furthermore, the distinct foci of satellite sequences seen at the beginning of cycle 14 are merged in an easily identifiable chromocenter in the apical part of the nucleus (Shermoen *et al.* 2010). Despite these differences, a connection between Rif1 and the replication program was still observed. At late anaphase of mitosis 14 and the onset of cycle 15, Rif1 was recruited rapidly to separating chromosomes, and appeared especially bright over the leading edge of the advancing chromosomes, the site of pericentric satellite sequences that constitute the bulk of the heterochromatin. As the telophase nucleus formed, bright Rif1 signal was seen over the compacted chromatin of the chromocenter before the recruitment of PCNA to the nucleus (Movie 4, Supplemental Figure 2). At the onset of S phase 15, PCNA was recruited only to euchromatic regions and it did not overlap Rif1 (Figure 2B). As S phase 15 progressed, PCNA was recruited to the edge of the compacted heterochromatin, where we have previously observed decompaction of HP1 staining chromatin (Shermoen *et al.* 2010). Although aggregated in a single mass in the nucleus (the chromocenter), individual satellite sequences remain as distinct subdomains and retain an individual replication schedule (Shermoen *et al.* 2010; Yuan and O’Farrell 2016). In agreement with this, Rif1 staining was progressively limited to more restricted regions within the chromocenter, with latest Rif1 foci dispersing toward the end of S phase (Figure 2B). Real time records again documented a close connection between the loss of Rif1 from chromatin and the recruitment of PCNA to the underlying region.

Having followed the progress of S phase globally, we next wanted to examine the replication of a specific heterochromatic sequence, the 1.686 satellite repeat. 1.686 is a simple repeated sequence at four loci, one to the left and one to right of the centromeres of both chromosomes 2 and 3. All these foci replicate together and show characteristic delays in their replication time in cycles 13 and 14. TALE-light probes can be injected into embryos and used to track the repetitive DNA in real time (Yuan et al., 2014; Yuan and O’Farrell, 2016). Purified mCherry labeled TALE-light protein recognizing the 1.686 repeat was injected into syncytial embryos and filmed. As reported previously, the TALE-light was gradually recruited to 1.686 during interphase, with the signal appearing as compact foci. During replication, the TALE-light signal became noticeably fuzzier, presumably due to the decompaction of the heterochromatic DNA during its active replication (Yuan *et al.* 2014). Upon completion of replication the TALE-light signal was again compact and obviously brighter. We use this reproducible behavior to indirectly follow the replication of 1.686. During cycle 13, we observed that Rif1 was recruited to 1.686 at the beginning of S phase, and disappeared immediately before the decompaction and replication of the repeat. We observed intermediate intensities of Rif1 signal on 1.686 in the minute preceding its decompaction. Rif1 dissociation was relatively rapid, but progressive with the Rif1 signal decaying over the minute preceding 1.686 decompaction. After completing replication, 1.686 lacked Rif1 for the remainder of the cycle (Figure 2C, Movie 5). Thus, Rif1 dissociates from this specific satellite immediately prior to the onset of its replication. We also examined fixed embryos using FISH probes to localize another heterochromatic repeat, AATAC, and showed that Rif1 was bound to this repeat AATAC during early S phase 14, but no such signal was observed in embryos aged to later in S phase after the replication of this sequence (Figure 2D). These observations show that Rif1 dissociates from specific satellite sequences upon their replication.

Our observations reveal that the dynamics of Rif1 interaction with chromatin parallel changes in replication timing. Rif1 association to satellite sequences began when these sequences first showed a slight delay in replication. While association was transient in the earlier cycles, Rif1 associated more persistently with satellites during the much-extended S phase of cycle 14. Most dramatically, Rif1 dissociated from individual satellite sequences at distinct times that align with the onset of PCNA recruitment to those sequences. The disappearance of the last foci of Rif1 staining marked the onset of replication of the latest replicating sequence and anticipates the end of S phase. These parallels suggest that binding of Rif1 to sequences might defer their replication until its dissociation. If so, analysis of the interaction may give us insights into the regulation of late replication and S phase prolongation.

### Steps in the appearance of localized Rif1 in foci

Late replicating sequences have to be specified before the onset of replication if they are to avoid early firing. In yeast and mammalian cells in culture, this is thought to occur well before replication at a critical time during G1, known as the timing decision point (Raghuraman *et al.* 1997; Dimitrova and Gilbert 2000). However in the G1-less embryonic cell cycles, the preparation for S phase is compressed and overlaps mitosis. Since cyclin:Cdk1 activity inhibits preparation of origins for replication, all of the preparations for replication happen between the down regulation of cyclin:Cdk1 at the metaphase anaphase transition and the onset of S phase upon entry into the next interphase. Real time observation of GFP tagged Orc2 protein revealed that Origin Recognition Complex (ORC) binds to the separating anaphase chromosomes prior to the midpoint of their separation (Baldinger and Gossen 2008). The MCM helicase is loaded shortly later in anaphase (Su and O’Farrell 1997). Replication begins at mitotic exit without obvious delay. This suggests that at the time of the transition from mitosis to interphase, there is already some type of feature that distinguishes the satellites so that they do not begin replication immediately (Shermoen *et al.* 2010). If Rif1 is to delay the replication time of specific sequences, we expect its recruitment to chromatin to occur during this period. Indeed, we observed binding of Rif1 to separating anaphase chromosomes before the recruitment of PCNA to the nucleus, which marks the start of interphase (Supplemental Figure 2B). However, as described below in cycles prior to cycle 15, the specificity of Rif1 binding to satellite sequences emerges in concert with progression into interphase.

We examined the initial binding of Rif1 by filming mitosis 13 and early interphase 14. We observed an abrupt onset of faint and generalized binding of Rif1 to the chromosomes during late anaphase/telophase as previously observed in fixed samples (Figure 3A) (Xu and Blackburn 2004; Sreesankar *et al.* 2015). Rif1 accumulation continued as the telophase nucleus formed. Although little was resolved in the compact and brightly staining telophase nucleus, when replication starts, rapid swelling of the interphase nucleus improved visualization. By about 1 minute into S phase 14, brighter foci of staining became clear, and much of the nuclear signal quickly declined except in foci clustered at the apex of the nucleus where the foci of pericentric satellite sequences lie (Figure 3A). Thus, while bound early, specificity in the binding became evident over a few minutes. In contrast, in the transition to cycle 15, the binding of Rif1 in anaphase was not uniform. During late anaphase 14, Rif1 was clearly enriched at the leading region of the chromosome mass, where the pericentric heterochromatin resides (Figure 3B, Movie 4). This localization to the heterochromatic region was visualized without interruption during the progression into interphase 15 and onset of replication. Thus, the character of anaphase binding of Rif1 changes between anaphase 13 and anaphase 14. Satellite sequences acquire some of the markings of heterochromatin such as methylation and HP1 localization during interphase of cycle 14. These heterochromatic marks might then guide the initial interaction of Rif1 at anaphase 14, but they would not be available to do so in anaphase 13.

**Figure 3.**
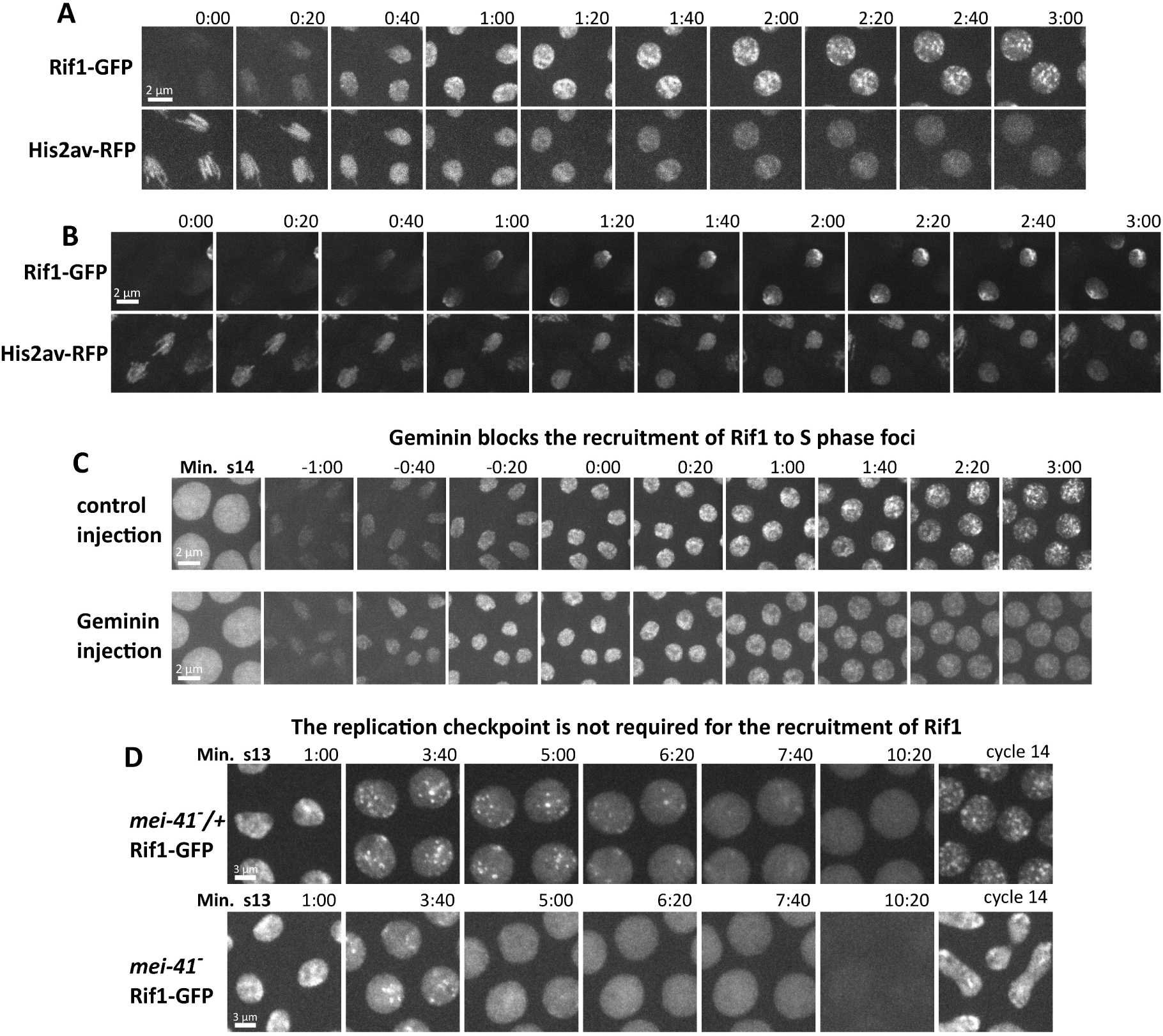
Two stages of Rif1 recruitment, their contributions to specificity, and reliance on origin licensing and the replication checkpoint. Time-lapse confocal microscopy on Rif1-GFP, His2av-RFP embryos showing the initial binding of Rif1 during transit from one cell cycle to the next. **(A)** During late anaphase 13, on the approach to cycle 14, chromosomes first exhibit faint and ubiquitous Rif1 staining that accumulates for about 1 min. During the following min as S phase begins, accumulation continues but is now localized to foci that become clearer as the nuclei swell. **(B)** During late anaphase 14, on the approach to cycle 15, the early stage of Rif1 staining shows some specificity for the pericentric regions of the chromosomes and forming chromocenter. This specificity is amplified as the interphase-15 nucleus forms and S phase begins. **(C)** Time-lapse imaging of Rif1-GFP during the normal transition from cycle 13 to 14 and in an embryo injected with purified geminin protein in interphase of 13. Times are indicated with reference to the start of S phase 14. The geminin block to pre-RC formation prevented the recruitment of Rif1 to foci, but did not block the initial generalized binding during mitotic exit. **(D)** Imaging of Rif1-GFP during cell cycle 13 in control (*mei41/+*) and *mei41* null embryos. The Mei41-dependent replication checkpoint is essential during cycle 13 to prevent premature entry into mitosis (10:20 frame). Rif1 foci still form in the absence of a replication checkpoint, but the Rif1 foci are lost earlier and the premature mitosis leads to bridging and defective cycle 14 nuclei.

Because recruitment of Rif1 began late in anaphase, just after the time when Orc2 was recruited to chromosomes, we wondered if pre-RC formation might be involved in Rif1 binding. Indeed prior work in a *Xenopus* extract system suggests that this is true (Kumar *et al.* 2012). To test this, we examined Rif1 recruitment after the injection of embryos with purified geminin protein during mitosis 13. Geminin blocks the formation of the pre-RC by inhibiting Cdt1, a key helicase-loading factor and prevents subsequent replication. Injection of geminin did not block the initial generalized binding of Rif1 to the late anaphase chromosomes or its nuclear accumulation in telophase nuclei, but did block the emergence of localized Rif1 foci (Figure 3C). In the absence of pre-RCs, Rif1 was diffusely localized throughout the nucleus during interphase 14.

### Recruitment of Rif1 to satellite sequences prior to the MBT does not require the replication checkpoint activity

Beginning in cycle 11, the gradual lengthening of interphase depends in part on the conserved DNA replication checkpoint (Sibon *et al.* 1997). During cycle 13, the checkpoint is required to delay activation of Cdk1 during S phase and prevent premature entry into mitosis. Embryos mutant for the checkpoint kinase ATR *(mei41)* fail to delay mitosis 13 sufficiently and so enter a catastrophic mitosis before the completion of replication resulting in massive chromosome bridging. Because Rif1 foci first became evident during these gradually slowing cycles, we wondered if the replication checkpoint might impact Rif1 recruitment. However, in *mei41* embryos, both the initial binding of Rif1 during anaphase of M12, and its subsequent recruitment to nuclear foci in interphase 13 were indistinguishable from control embryos (Figure 3D). We did observe that the disappearance of Rif1 foci was accelerated in *mei41* embryos, and Rif1 was lost from chromatin before entry into a catastrophic mitosis 13. We conclude that the recruitment of Rif1 to late replicating sequences is independent of the replication checkpoint, but that timing of Rif1 dispersal is accelerated in its absence.

### Cdk1 activity promotes Rif1 release from chromatin during S phase

Prior work demonstrated that down regulation of Cdk1 activity at the MBT plays a key role in extending S phase (Farrell et al, 2012; Farrell and O’Farrell, 2013; Yuan et al., 2016). Cdk1 activity during the earlier rapid cell cycles was shown to be required for satellite sequences to replicate early, and to sustain short S phases. The gradual decline in mitotic activators during cycles 11-13 contributes to the progressive lengthening of S phase. Finally at the MBT, the abrupt drop in Cdk1 activity is required for late replication and the resulting prolongation of S phase. These changes to Cdk1 parallel the changes we observed for Rif1 localization, so we investigated the connection between them.

Experimental reduction of Cdk1 activity by injection of RNAi against the 3 mitotic cyclins (A, B, and B3) during cycle 10 arrests embryos in interphase 13, and extends the length of S phase 13 from 13 minutes to an average of 19 minutes (Farrell et al., 2012). Cyclin knockdown also affected the dynamics of Rif1 chromatin association. In arrested embryos, foci of Rif1 binding persisted for an average of 16 minutes, compared to 8 minutes in control-injected embryos (Figure 4A, 4C). This delay in the loss of Rif1 foci paralleled, and slightly preceded, the delayed appearance of late replicating PCNA signal. In contrast, as previously reported, increasing Cdk1 activity during cycle 14, by injecting mRNA for the mitotic activator Cdc25, shortened S phase from over an hour to an average of 22 minutes. Experimental Cdk1 activation also changed the Rif1 program. In Cdc25 injected embryos, foci of Rif1 association were evident for only an average of 18 minutes compared to 66 minutes in control-injected embryos (Figure 4B, 4C). The accelerated loss of Rif1 foci slightly preceded the advanced appearance of late replicating PCNA foci (Movie 6). Both experiments demonstrate that the timing of Rif1 dissociation from chromatin and the subsequent initiation of late replication are sensitive to the activity of Cdk1 during S phase.

**Figure 4.**
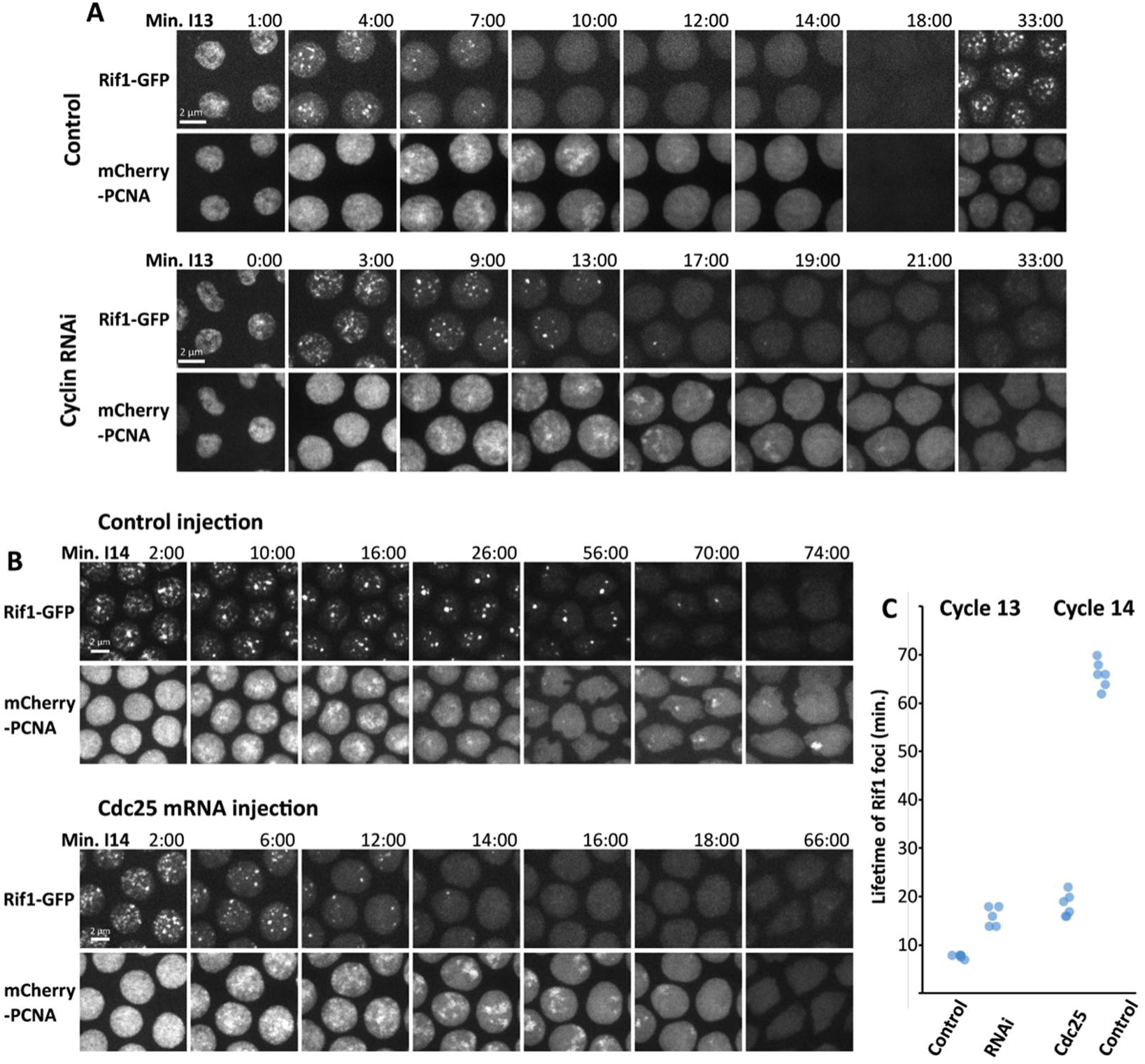
Manipulation of Cdk1 activity alters the lifetime of Rif1 foci and the duration of S phase. **(A)** Stills from time-lapse imaging of Rif1-GFP and mCherry-PCNA during cycle 13 after injection of either buffer (control) or dsRNA against the three mitotic cyclins (A, B, and B3) in cycle 10. The knockdown of the cyclins increases the persistence of Rif1 foci, extends S phase (PCNA staining) and blocks progression to cycle 14. The time of the last frame showing a Rif1 focus is indicated as the lifetime of Rif1 foci in (C). (B) Stills from time-lapse imaging of Rif1-GFP and mCherry-PCNA during cycle 14 after injection of either buffer (control) or mRNA encoding the Cdk1 activator Cdc25 (twine) in cycle 13. Artificial activation of Cdk1 during S phase 14 accelerates both the dissociation of Rif1 and late replication. In the control embryo, Rif1 foci were detected 70 min into S phase 14, and prominent foci of PCNA were documented at 74 min. Expression of Cdc25 resulted in the loss of Rif1 foci by 16:00 and the final late replicating PCNA signal documented was at 18:00 (Movie 6). **(C)** Plot of lifetime of Rif1 foci in minutes during either S phase 13 or S phase 14 following the indicated injection.

These results show that Cdk1 activity in early cycles normally promotes Rif1 dissociation from chromatin in pre-MBT embryos and that an artificial increase in Cdk1 in cycle 14 can do the same. The parallel effects of the experimental manipulations on Rif1 association and the progress of S phase further support suggestions that Rif1 association suppresses replication and that activation of origins in satellite sequences occurs in conjunction with Rif1 dissociation. The influence of Cdk1 on Rif1 suggests that the developmental program of Cdk1 down regulation guides the observed changes in Rif1 dynamics and thereby, S phase duration. In light of these findings we can interpret the accelerated loss of Rif1 in *mei41* embryos (Figure 3D) as a consequence of the faster activation of S phase Cdk1 in the absence of the replication checkpoint (Deneke *et al.* 2016). The observations suggest that activity of the Cdk1 kinase directly or indirectly regulates Rif1 interaction with chromatin.

### A phospho-site mutant Rif1 does not release from heterochromatin and prevents completion of DNA replication

Work from both *S. cerevisiae* and *S. pombe* indicates that the kinases CDK and DDK act on conserved phosphorylation sites to inhibit Rif1 and limit its ability to inhibit replication (Hiraga et al., 2014; Dave et al., 2014). In these yeasts, phosphorylation of Rif1 is thought to block Rif1’s ability to recruit PP1, hence preventing Rif1 inhibition of replication initiation. Like yeast Rif1, the dipteran Rif1 homologs have a cluster of conserved CDK and DDK sites near the phosphatase interaction motif in the C-terminal part of the protein (Figure 5A, Supplemental Figure 5). This region of Rif1 is also proposed to contain a DNA binding domain (Sreesankar *et al.* 2012; Zimmermann *et al.* 2013). Interestingly these potential phosphorylation sites cluster in a region of the protein predicted to be of high intrinsic disorder (Supplemental Figure 5) a feature associated with phosphorylation events that regulate interactions (Holt *et al.* 2009). We further examined the role of the more C-terminal potential phosphorylation sites in regulating the late replication activity of Rif1.

**Figure 5.**
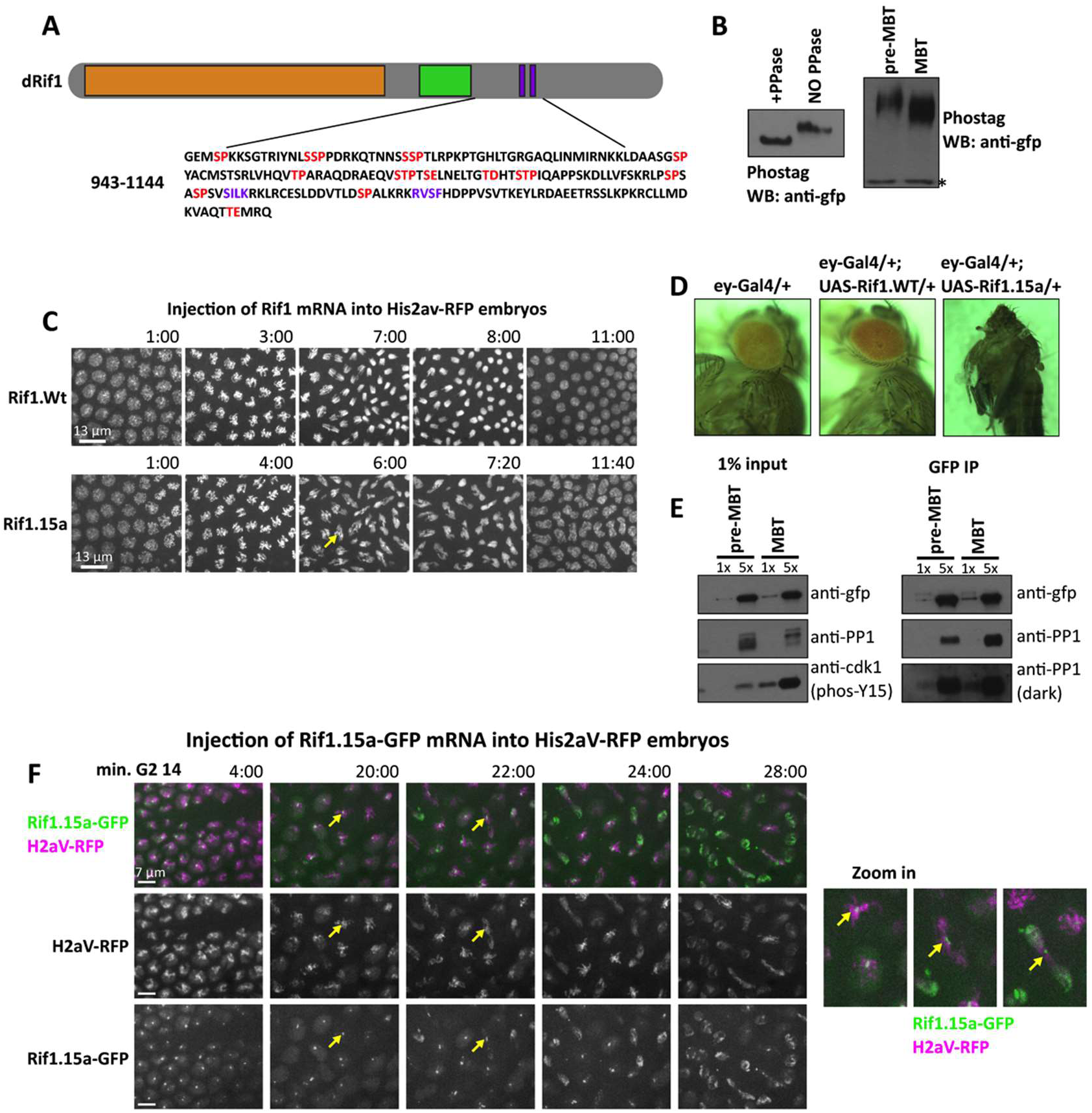
Phospho-site mutant Rif1 does not dissociate from chromatin and blocks replication. **(A)** Schematic (above) of Rif1 protein with N-terminal HEAT repeats (orange), putative DNA binding domain (green), and PP1 interaction motifs (purple boxes). The amino acid sequence (below) of the indicated portion of the Rif1 protein shows candidate CDK and DDK phosphorylation sites (red). These 15 S/T residues were mutated to alanine to create the Rif1.15a phospho-mutant allele. **(B)** Left panel shows anti-gfp Western blots used to detect Rif1-GFP from 1-hour old embryos. Protein extracts were treated with lambda phosphatase or buffer only for 1 hour and then were run on SDS-PAGE gels cast with phostag to decrease the migration of phosphorylated proteins. Right panel shows phostag western blots used to detect Rif1-GFP from embryos aged to one hour (pre-MBT) or to 2.5 hours (MBT). Asterisk denotes background band. **(C)** Ectopic expression of the indicated versions of Rif1 by injection of mRNA during cycle 11. After injection, embryos were filmed during mitosis 13. The yellow arrow points to an example of an anaphase bridge. **(D)** Images of flies (left) and a pharate pupa (right) show the consequence of expression of either Rif1 (middle) or Rif1.15a (right) in the developing fly eye/head capsule using Eyeless-Gal4. **(E)** Anti-GFP immunoprecipitation followed by Western blotting with the indicated antibodies. Protein extracts were prepared from embryos harvested before the MBT (1 hour old) or after aging to the MBT (2.5 hours old). Phos-Y15 (left-bottom) indicates the inactive phosphorylated form of Cdk1. **(F)** Frames from live imaging of nuclei starting in G2 of 14 and progressing into a defective mitosis following expression of Rif1.15a-GFP from injected mRNA. Yellow arrows indicate extensive anaphase bridging with a residue of Rif1.15a-GFP. Magnified images (right) show progression of one nucleus.

To determine if Rif1 is phosphorylated *in vivo,* we examined Rif1 from 1-hour old embryos, a time when the abundant maternally provided Rif1 protein would be held inactive by Cdk1. Protein extracts from pooled staged embryos were run on a phos-tag SDS-PAGE gel before or after treatment with lambda phosphatase, and subjected to western blotting. All detectable Rif1 exhibited a phosphatase-reversed shift, indicating that most Rif1 is phosphorylated in the pre-MBT embryo (Figure 5B). Comparison of protein samples from pre-MBT and MBT age embryos revealed that the extent of the shift is somewhat reduced at the MBT stage, consistent with developmentally regulated reduction in the degree of phosphorylation (Figure 5B).

Two studies in yeast suggested that phosphorylation of Rif1 inhibits its function by blocking interaction with PP1a (Dave et al., 2014 and Hiraga et al., 2014). *Drosophila* Rif1 has previously been shown to bind to PP1a, so we looked for a change to this interaction in conjunction with the developmental down regulation of Cdk1. As shown above, high Cdk1 activity in early pre-MBT embryos suppresses Rif1 association with chromatin and promotes a short S phase, whereas down regulation of Cdk1 at the MBT is required for Rif1 mediated extension of S phase. Rif1 was immunoprecipitated out of lysates derived from 20-minute collections of embryos aged for either 30 minutes (pre-MBT) or for 2 hours and 15 minutes (post-MBT). While we expected minimal association of PP1 to Rif1 in pre-MBT extract when Rif1 is robustly phosphorylated, western blotting showed a robust PP1 signal (Figure 5E). Additionally, Rif1 immunoprecipitated from the post-MBT extract was accompanied by a similar amount of PP1. Hence, the interaction between Rif1 and PP1a, at least at bulk levels, is not regulated in a way that explains the onset of late replication, and Rif1 appears to interact effectively with PP1 at early stages when it is phosphorylated and held inactive by Cdk1 kinase.

To further explore possible regulation of Rif1 by phosphorylation, we mutated candidate phospho-sites. We selected 15 S/T residues within CDK or DDK consensus motifs located in the C-terminus of Rif1 and mutated them to alanine (Figure 5A) to prevent their phosphorylation. We reasoned that this phospho-site mutant Rif1, hereafter referred to as Rif1.15a, might not be inhibited by S phase kinases and so act as a dominant gain-of-function allele. Ectopic expression of Rif1.15a during the blastoderm divisions by injection of in vitro transcribed mRNA causes extensive anaphase bridging as typically seen when DNA replication is incomplete (Figure 5C). We never observed this effect after injection of mRNA encoding wild-type Rif1 arguing that this is not a simple overproduction phenotype. In addition, we generated transgenes expressing either wild-type Rif1 or Rif1.15a under UAS control. Overexpression of Rif1 in the eye imaginal disc (and more weakly throughout much of the head capsule) using the eyeless-Gal4 driver did not disturb eye formation whereas expression of Rif1.15a caused complete lethality with pupae developing into headless nearly adult flies (Figure 5D). The severity of this phenotype suggests that expression of Rif1.15a disrupted the early proliferative period of the eye-antennal disc. These assays suggest that Rif1.15a has a damaging gain-of-function action, as might be expected if it were immune to regulation of its ability to inhibit DNA replication.

Because our data indicated that the chromatin binding of Rif1 was regulated during S phase by Cdk1 activity, we examined the influence of mutation of the phospho-sites on Rif1 localization. Using mRNA injection, we expressed a GFP tagged version of Rif1.15a and followed its localization and the consequence of its expression in live records. Embryos were injected with Rif1.15a-GFP mRNA during cycle 12. The timing of this injection allowed sufficient fluorescent protein to accumulate that its behavior could be followed throughout cell cycle 14, while still minimizing the damage due to induced catastrophic mitoses in the earlier cycles. Rif1.15a was recruited to specific chromatin foci normally at the start of S phase, indicating that the mutated residues are not required for the binding specificity of Rif1. Imaging Rif1.15a-GFP throughout cycle 14 yielded several interesting results. In contrast to the dynamics of the wild-type protein, Rif1.15a continued to accumulate on heterochromatin throughout most of S phase 14. It then showed a slow decline but remained bound in the following G2 and into mitosis 14 (Figure 5E). The Rif1.15a foci on newly condensed mitotic chromosomes were localized to pericentric regions, where the satellites reside. On progression into anaphase, bridges were seen connecting the separating chromosomes and Rif1.15a-GFP specifically labeled these chromatin bridges (Figure 5E inset). As bridged nuclei exited mitosis, an unbound pool of Rif1.15a was recruited to the chromocenter, presumably after new pre-RCs were loaded. This abrupt recruitment of Rif1.15a shows that mutation of the selected sites did not fully eliminate cell cycle regulated behavior of Rif1. Nonetheless, the dramatic consequence of the phosopho-site mutations shows that these sites contribute importantly to Rif1 dissociation from chromatin and that in the absence of dissociation Rif1.15a is capable of blocking replication to give anaphase bridging when chromosomes are driven into mitosis.

### Rif1 is dispensable for survival and fertility

A recent study reported that Rif1 was an essential gene in *Drosophila* based on partial lethality of a ubiquitously expressed RNAi against Rif1 (Sreesankar *et al.* 2015). We created a precise deletion of the Rif1 ORF using Crispr-Cas9 (Supplemental 6). While the mutation does cause reduction in survival, we find that the homozygous *rif1* null gives viable, reproductively competent flies that can be propagated as a stock. Zygotic loss-of-function *rif1* mutants develop to adulthood with a reduced survival and a male to female ratio of 0.4 (n = 189 flies). Additionally, embryos laid by *rif1* mutant mothers (hereafter called *rif1* embryos) have a reduced hatch rate (55% of control, n = 500 embryos). We conclude that Rif1 is dispensable in *Drosophila* at least in our genetic background, and we suspect that the previously reported lethality/sterility included enhancement of the loss-of-function phenotype by off-target effects of the RNAi. Though surprised by the viability of the Rif1 delete, it provided an opportunity to examine the phenotype of complete absence of function.

### Rif1 is required for the onset of late replication and the prolongation of S phase at the MBT

Next we utilized the *rif1* null mutation to assess the role of Rif1 in the onset of late replication during cycle 14. As discussed previously, real time records of fluorescent PCNA allow us to follow the progress of S phase and estimate its overall length. In a wild-type S phase 14, PCNA is widely distributed throughout most of the nucleus during the first 10 minutes, but resolves into bright puncta that are obvious for over an hour into interphase, after which only a dim nuclear background is visible. During the final 30 minutes of S phase, a small number of bright PCNA foci appear and then disappear sequentially as a result of a protracted schedule of late replication (Figure 6A). By using the disappearance of the last PCNA focus as an indication of the completion of replication, we determined that the average S phase 14 lasts for 73 minutes (Figure 6A). In contrast, S phase 14 in *rif1* embryos from *rif1* mothers was significantly shorter, lasting an average of 27 minutes.

**Figure 6.**
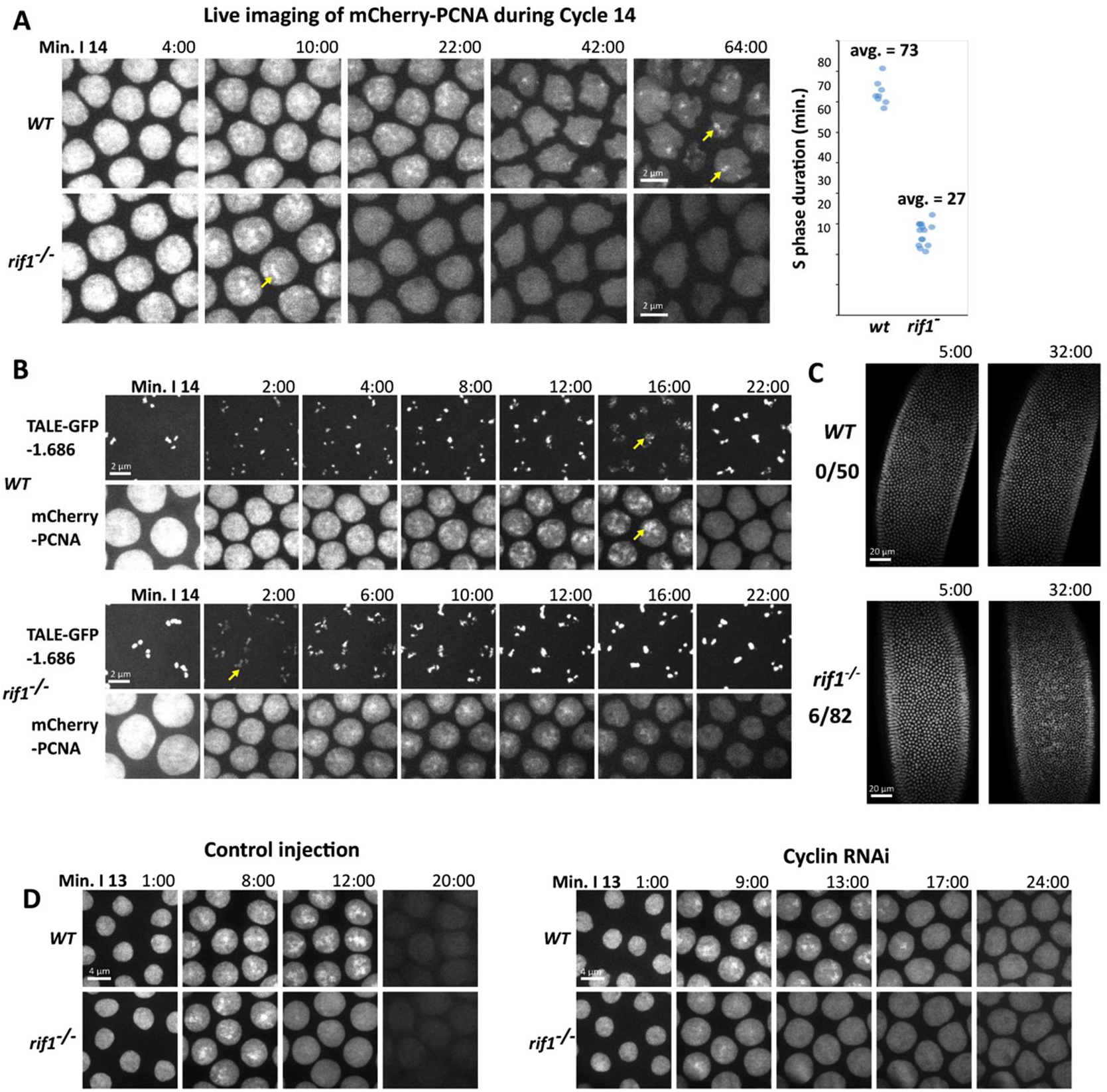
*rif1* is required for the prolongation of S phase at the MBT. **(A)** Stills from time-lapse imaging of mCherry-PCNA during S phase 14 in wild-type (top) or *rif1* mutant (bottom) embryos. Yellow arrows indicate late replication foci. The plot on the right displays the total duration of S phase 14 in wild-type and *rif1* embryos. Each data point represents a distinct embryo. The duration of S phase was scored from real time records of PCNA dynamics. **(B)** Stills from time-lapse imaging following the TAIL-light probe for the 1.686 satellite (upper) and mcherry-PCNA to follow replication during S phase 14 in wild-type and *rif1* embryos. Yellow arrows indicate the decompacted signal from the 1.686 TALE-light during the replication of the 1.686 repeat. **(C)** Wild-type (top) or *rif1* (bottom) embryos were injected with GFP-HP1a to visualize nuclei and filmed during cycle 14. 6/82 *rif1* embryos underwent an early mitosis 14 (Movie 7). **(D)** Still frames from movies following mCherry-PCNA during S phase 13 in wild-type and *rif1* embryos after either control or triple cyclin RNAi injection. Mitotic-cyclin knockdown arrests the cell cycle, but extends S phase in wild-type but not *rif1* embryos.

Additionally, there were clear differences in the appearance of the PCNA signal as S phase progressed. The initial period of widespread early replication resolved into PCNA foci, but appearance of these foci was more simultaneous than sequential, and there was no protracted sequence of late foci (Figure 6A). Thus, this S phase 14 resembled the earlier S phases, in showing a marginal extension with slightly delayed replication of some foci. However, the normally dramatic extension of S phase 14 is dependent on *rif1.* Failure to extend S phase 14 was observed in all embryos derived from homozygous mothers even though heterozygous fathers where included in the cross. We conclude that maternal Rif1 is required for the normal extension of S phase 14.

Because we had observed that the satellite 1.686 recruited Rif1 during S14, we wondered if its replication time would be altered in *rif1* embryos. To measure the replication time of this repeat, we injected the GFP-1.686 TALE probe into wild-type and *rif1* embryos expressing mcherry-PCNA and recorded S phase 14. We used the transient recruitment of mcherry-PCNA, and the obvious decompaction of the marked 1.686 sequences as indictors of active replication of 1.686. In control embryos, the 1.686 repeat began replication 18 minutes into S14 and completed by 30 minutes (Figure 6B). In contrast, in *rif1* embryos, 1.686 began replication by 4 minutes into S14 and completed by 13 minutes (Figure 6B). We conclude that the 1.686 satellite sequence replicates with a minimal delay in S phase 14 in the absence of Rif1, and that the normally substantial delay requires Rif1.

During the MBT, the embryo degrades both the mRNA and protein of the mitotic activator Cdc25. This allows the introduction of the first embryonic G2 after the completion of the prolonged S phase 14. Mitosis 13 is then the last synchronous division during development, and mitosis 14 relies on the developmentally patterned zygotic expression of new Cdc25. Perturbations that interfere with the downregulation of Cdk1 can lead to an additional synchronous mitosis. Additionally, since down regulation of Cdk1 occurs during S phase 14, the embryo can also progress to an additional synchronous mitosis if S phase is eliminated or dramatically shortened, as seen following injection of geminin or alpha-amanitin, respectively (McCleland *et al.,* 2009; Shermoen *et al.,* 2010). We noticed that a small number of *rif1* embryos (7%) executed an early mitosis 14 at approximately 30 minutes into interphase. The observed mitoses were complete, with no bridged chromosomes, but there was substantial nuclear fall in (Figure 6C, Movie 7). We interpret the incomplete penetrance of the extra-division phenotype to be an indication that the duration of S phase in *rif1* null embryos is close to a threshold, so that in most embryos the cyclin:Cdk1 activity declines enough to introduce a G2, but in some embryos the residual maternal cyclin:Cdk1 function remains high enough to trigger mitosis upon the completion of the shorten S phase. The result shows that Rif1 is important for reliable coordination of the MBT.

Previous work has shown that before the MBT, Cdk1 is required for driving a short S phase. Our finding that Cdk1 activity suppresses Rif1 function in cycle 14 (Figure 4) suggests an explanation for the early role of Cdk1 in promoting short S phases; interphase activity of Cdk1 prevents Rif1 from prematurely introducing late replication. This model predicts that reducing Cdk1 activity would not be able to prolong a pre-MBT S phase in the absence of *rif1.* To test this idea we examined S phase 13 in wild-type or *rif1* embryos after knockdown of mitotic cyclins, essential activators of Cdk1, by RNAi. Wild-type control injected embryos exhibit transient but obvious foci of PCNA staining during late S phase 13 and replicate their satellite sequences with a slight delay in this S phase. While *rif1* embryos still exhibit PCNA foci, these foci are even shorter lived (Figure 6D). Following injection of mitotic-cyclin RNAi, wild-type embryos increased the duration of S phase 13 from 14 minutes to 20 minutes. In contrast, *rif1* embryos did not extend S phase after cyclin knock down (Figure 6D). This demonstrates that the requirement for Cdk1 in the timely completion of S phase 13 can be bypassed by loss of *rif1*. However, S phase 13 in the *rif1* mutant embryos is still longer than very early embryonic S phases, which can be as short as 3.4 min. Thus, Cdk1 down regulation of Rif1 contributes to S phase prolongation in cycle 13, but there must be additional factors influencing the progressive prolongation of early cycles.

### *Cdc7* is essential during the early embryonic S phases and the requirement can be substantially bypassed by removal of *rif1*

The Cdc7-Dbf4 kinase complex (or DDK – Dbf4 dependent kinase) is required for origin initiation in many systems. In *S. cerevisiae,* the essential function of DDK is the phosphorylation of the MCM helicase complex during pre-RC activation (Sheu and Stillman 2010). In both *S. pombe* and *S. cerevisiae,* the deletion of *rif1* partially rescues S phase and viability in *cdc7* mutants. Two interactions have been proposed to contribute to this finding. DDK appears to phosphorylate and inactivate Rif1, thereby activating replication by a derepression input. This input would be dispensable in a *rif1* mutant. Additionally, Rif1 is thought to inhibit replication by recruiting PP1 to the origin and dephosphorylating the MCM helicase, an action that opposes DDK. The loss of this opposing activity in a *rif1* mutant would reduce the required activity for MCM phosphorylation that might then be provided by residual DDK function, or an alternative kinase such as a Cdk. We wanted to assess the possible parallels in *Drosophila* to clarify the involvement of Rif1 in replication control.

Despite strong conservation, *cdc7* has not been well studied in *Drosophila.* In *Drosophila, cdc7* is an essential gene, and recent work has demonstrated that in complex with the *dbf4* ortholog *chiffon, Drosophila* Cdc7 can phosphorylate Mcm2 *in vitro,* and that *cdc7* is required for endocycle S phases in follicle cells (Stephenson *et al.* 2014). However, the function of DDK during mitotic S phase has not been described.

To address this issue we first tagged endogenous *cdc7* with GFP using CRISPR-Cas9. The resulting *cdc7-GFP* stock was healthy and fertile indicating that the tag did not disrupt the essential function of Cdc7. Time-lapse imaging of Cdc7-GFP embryos during syncytial development revealed that Cdc7 localization was cell cycle regulated. Cdc7 was nuclear during interphase, dispersed into the cytosol during mitosis, and was rapidly recruited to late anaphase chromosomes and concentrated in the telophase nucleus (Figure 7A). Initial Cdc7 recruitment featured two transient bright foci followed by fine puncta suggesting that it is recruited to chromatin at the time at which pre-RCs are undergoing activation during the syncytial cell cycles. Comparison of nuclear Cdc7 at equivalent times in subsequent cell cycles (early S phase) leading up to the MBT revealed a decline in the per-nucleus protein level, suggesting that titration by increasing number of nuclei and/or declining protein levels result in diminishing availability of Cdc7 to fire origins in later cycles (Figure 7B, Movie 8).

**Figure 7.**
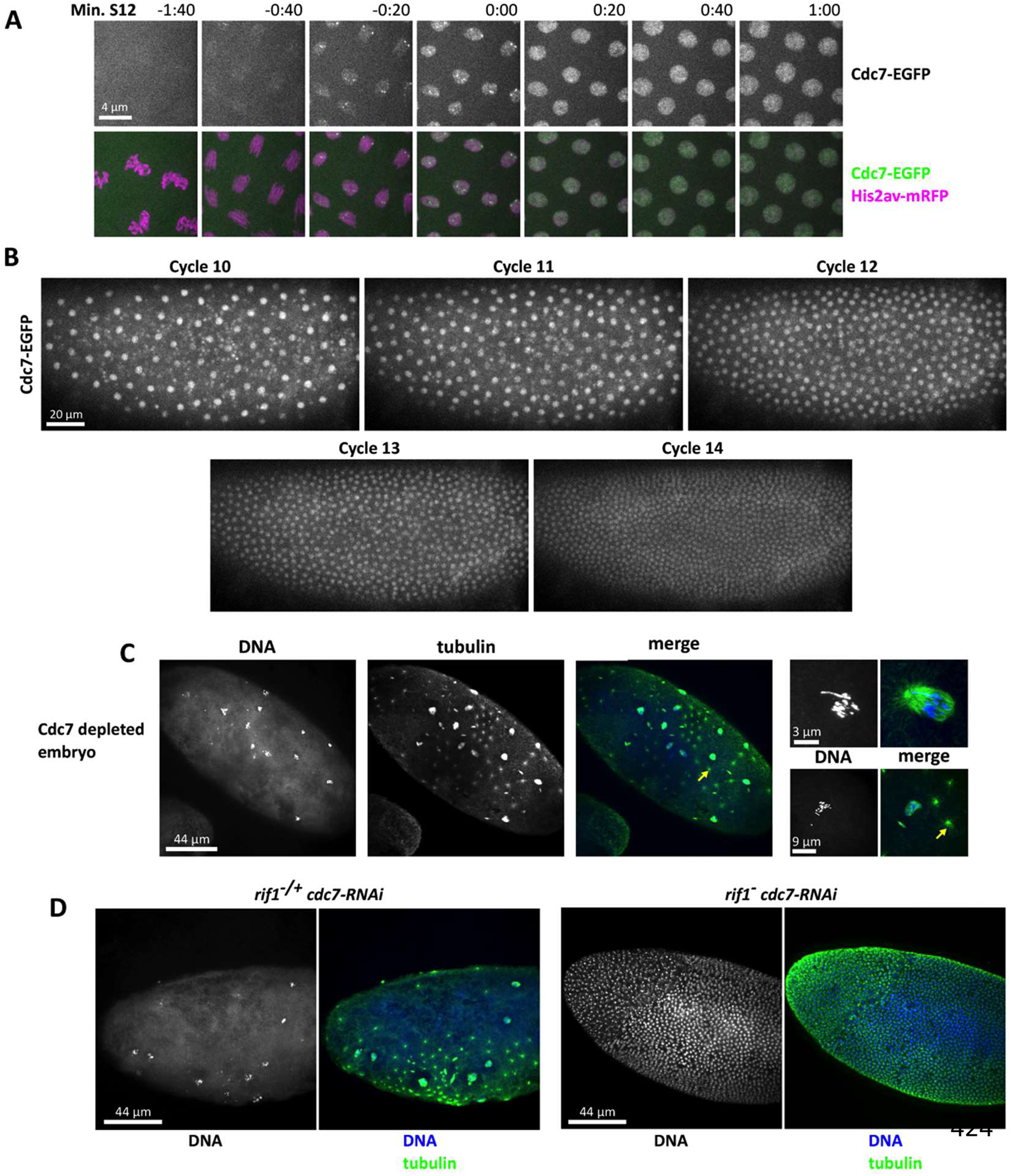
Deletion of *rif1* rescues cell cycles to Cdc7 depleted embryos. **(A)** Cell cycle regulated recruitment of Cdc7 to the nucleus during early interphase shown by time-lapse microscopy of Cdc7-GFP, His2av-RFP during nuclear cycle 12 using a 100x objective. **(B)** Live imaging of Cdc7-GFP embryos during cycles 10-14 reveals dilution of nuclear Cdc7 over the pre-MBT divisions. (C) Fixed cdc7 depleted embryos stained for DNA and tubulin. Yellow arrow indicates a centrosome not associated with a DNA mass. Adjacent enlarged images display magnified views of abnormal mitotic structures. **(D)** Fixed *rif1* mutant embryo after maternal RNAi against cdc7 stained for DNA and tubulin. Deletion of *rif1* restores cell cycles and early development to cdc7 depleted embryos.

To determine if *cdc7* is required for DNA replication during the early cell cycles, we depleted DDK from the embryo using maternal-tubulin Gal4 to drive RNAi against *cdc7* during oogenesis. This setup did not interfere with egg production, although we found that expression of RNAi against *cdc7* using the earlier acting MTD-Gal4 did cause severe defects in egg morphology, indicating that *cdc7* does play a role in the female germline. Embryos depleted of Cdc7 using maternal-tubulin Gal4 failed to hatch. Cytological examination of Cdc7 depleted embryos revealed penetrant defects during the early preblastoderm divisions. We observed multiple highly fragmented DNA masses that were unevenly distributed in the embryo interior, along with scattered abnormal mitotic structures (Figure 7C). In all cases nuclei failed to form a blastoderm, although in many cases centrosomes appeared to continue to duplicate in the absence of any associated DNA (Figure 7C). Such a dissociation of the embryonic nuclear and centrosome cycles has been described before in embryos injected with the DNA polymerase inhibitor aphidocolin during the pre-blastoderm cycles (Raff and Glover 1988). We conclude that Cdc7 is essential for effective nuclear cycles in the early embryo consistent with a requirement in DNA replication.

Next we tested for a genetic interaction by depleting Cdc7 from *rif1* mutant embryos. In the absence of *rif1,* 2% of Cdc7-depleted embryos hatched. While the Cdc7-depleted embryos from mothers heterozygous for *rif1* (*rif1+*) never hatched. Cytological examination demonstrated that removal of *rif1* restored cell cycle progression to Cdc7-depleted embryos. *rif1* mutants completed substantially more cell cycles than heterozygous control embryos, with most progressing to the blastoderm stage. Many of the blastoderm rescued embryos showed abnormalities such as substantial nuclear fall in, disorganized nuclear spacing, and non-uniform nuclear density. Nonetheless, some of the observed embryos attempted cellularization and gastrulation (Figure 7D) with the rare cases of hatching indicating occasional success. We conclude that deletion of *rif1* can restore embryonic cell cycle progression and early development to Cdc7-depleted embryos. However, this rescue is incomplete, likely because of altered regulation of the restored S phase in such doubly defective embryos.

### The Rif1-regulated onset of late-replication precedes the establishment of heterochromatic marks

Early embryonic chromatin lacks specializations that come to distinguish different regions of the genome at later stages. Thus, the order of appearance of different specializations can give us insight into the hierarchy of regulation. Since late replication is considered a feature of heterochromatin, we expected its emergence to follow the embryonic appearance of the hallmarks of heterochromatin. Our recent work described the onset of localized H3K9me2/3 and HP1a (Yuan and O’Farrell 2016). While a very low level of modification and chromatin bound HP1a was detected before the MBT, a period of abrupt accumulation of HP1a and more extensive H3K9me2/3 modification only occurred later, during S phase 14. Here we explore the relationship between this HP1a-centered program and the action of Rif1 in the control of late replication.

During cycle 14, Rif1-GFP and HP1a-RFP bound to localized foci in the same position, but never at the same time. Rif1 foci disappear sequentially during cycle 14, while HP1a is diffusely localized in early cycle 14 and is subsequently recruited to apical foci that multiply and intensify. Importantly, in live imaging of Rif1-GFP and HP1-RFP, we don’t observe any overlap between the two signals in S14 (Figure 8A). This finding is in accord with our finding that PCNA foci and Rif1 do not overlap (above), and our previous demonstration that PCNA and HP1 foci do not overlap in cycle 14 (Shermoen *et al.,* 2010). Together with the timing of recruitment of these proteins to specific satellites, these observations show a sequence in which satellites lose associated Rif1 before they initiate replication as marked by PCNA, and then complete replication before binding HP1. We conclude that the introduction of late replication by Rif1 precedes the binding of HP1a in cycle 14. In contrast, at the start of cycle 15 both Rif1 and HP1a are rapidly recruited to the chromocenter, with Rif1 binding slightly earlier (Figure 2B, 3B), and we have previously detected an influence of HP1 on replication timing in this cycle (Yuan and O’Farrell, 2016).

**Figure 8.**
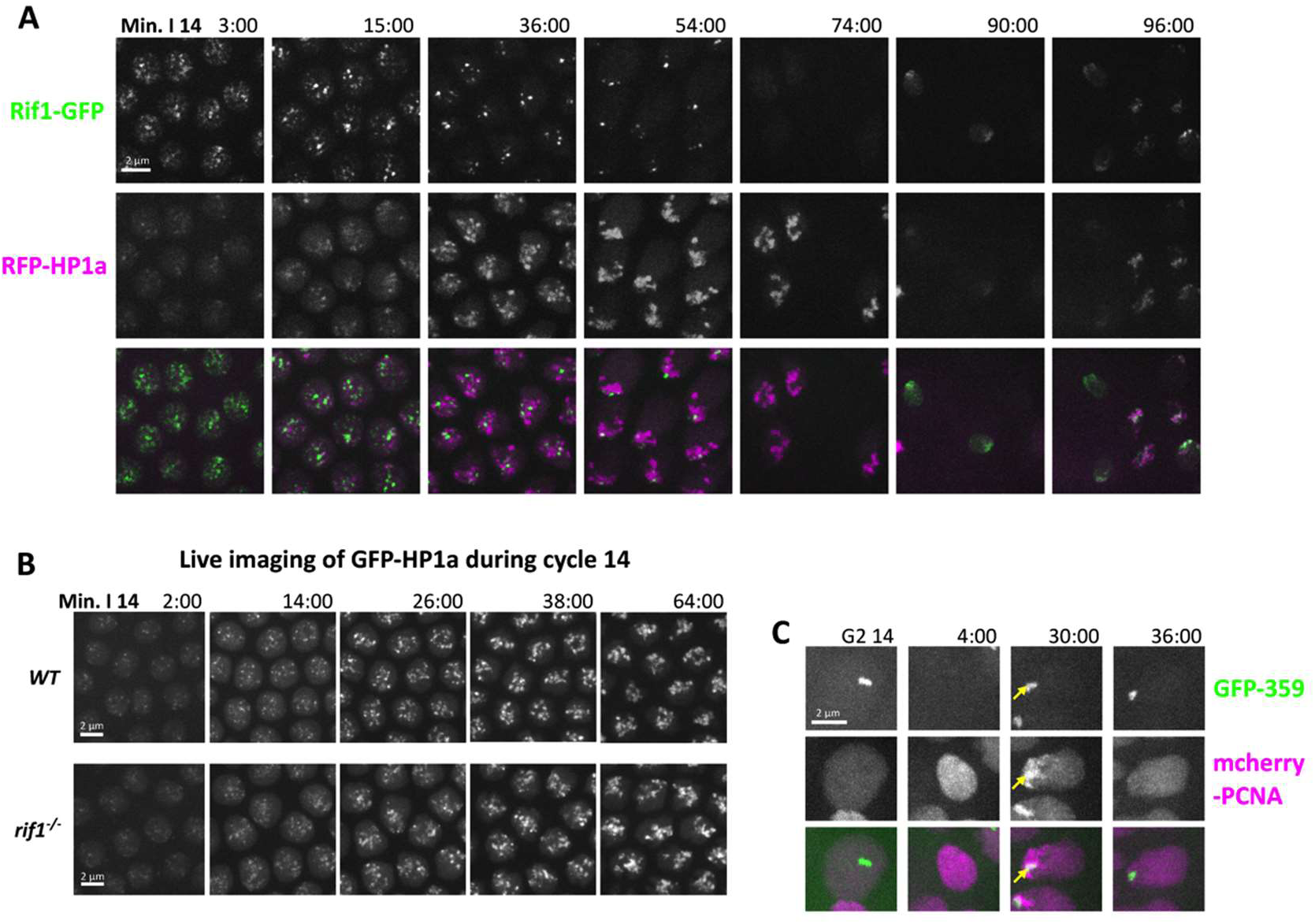
Sequential appearance at satellite sequences suggests that Rif1 influences late replication prior to an impact of HP1a. **(A)** Still images from time-lapse imaging of Rif1-GFP and RFP-HP1a during cycle 14 (early S phase through mitosis). During S phase the number of Rif1 foci decline as the number of HP1a foci increase. G2 nuclei lack Rif1 foci, but retain strong HP1a foci (74 min). During the asynchronous mitosis 14 (74/90/96 min), both proteins are lost, but are rapidly recruited to late anaphase chromosomes (90/96 min). **(B)** Still images from time-lapse imaging of GFP-HP1a protein injected into wildtype embryos (above) and *rif1* embryos (below) during S phase 14. HP1a recruitment to the heterochromatin proceeded similarly in control and mutant embryos. **(C)** Still images from timelapse imaging of the replication of the 359-satellite progressing from G2 of cycle 14, until completion of its replication in cycle 15. Note that the TAIL-light signal is not immediately visible following mitosis (4:00 min frame). We previously described how HP1a binds to the 359 bp repeat following its replication during S phase 14 and subsequently delays its replication in S phase 15. In *rif1* embryos, the 359-repeat also replicated late in S phase 15 (In frame 30:00, yellow arrow indicates where the 359 TALE signal overlapped with PCNA).

Given the earlier binding of Rif1 to satellite sequences in cycle 14, Rif1 might direct subsequent heterochromatin formation. In fission yeast, Rif1 is required for the maintenance of silencing at some heterochromatic sites in the genome (Zofall *et al.* 2016). We examined whether Rif1 influences the emergence of HP1a-bound heterochromatin. GFP-HP1a was injected into either wild-type or *rif1* null embryos and imaged during cycle 14. We observed no difference between HP1a recruitment between the two genotypes, indicating that HP1a binds independently of Rif1 in the fly embryo (Figure 8B).

We had previously found that the recruitment of HP1a to the 359 bp-repeat satellite-sequence during cycle 14 occurred only after its replication and was unimportant to the replication timing of this satellite in S phase 14. However, this recruitment of HP1a was required for a shift of 359 replication to a much later time in S phase of cycle 15 (Yuan and O’Farrell 2016). Because our results demonstrate that the recruitment of HP1a to the heterochromatin during S phase 14 is independent of Rif1, we wondered if the replication time of 359 was altered in *rif1* embryos in cycle 15. Live imaging of mCherry-PCNA expressing *rif1* embryos injected with GFP-359 TALE probes indicated that 359 still replicates late during S15 (Figure 8C). We conclude that an HP1a dependent program can delay replication of 359 sequences in cycle 15 without Rif1 input. It seems likely that this HP1a dependent program operates to influence the replication of many heterochromatic sequences after cell cycle 14.

## DISCUSSION

We have combined fly genetics with live imaging to reveal and study a novel developmental role of *Drosophila* Rif1 in controlling the onset of late replication during embryogenesis. The findings suggest how a conserved feature of constitutive heterochromatin, its late replication, first arises during embryogenesis. The dominant role that Rif1 plays in late replication at this developmental stage gave us an unobstructed view of this function. We show that chromatin association of Rif1 during development is coordinated with onset of its action in delaying replication, and that Rif1 dissociation from regions of chromatin during S phase is temporally coupled to onset of the delayed replication of affected sequences. In addition, the developmental context of our study allows ordering of the events that introduce different features to the heterochromatin.

We show that Cdk1 activity promotes dissociation of Rif1 from chromatin and inhibits its function. Importantly, the ability of Cdk1 to limit Rif1 function provides a key link in the developmental control of the MBT. A developmental decline in Cdk1 triggers the extension of the cell cycle at the MBT (Farrell *et al.,* 2012; Yuan *et al.,* 2016). However, it was not previously apparent how a decline in Cdk1, a mitotic kinase, would trigger the abrupt extension of S phase that marks the onset of cell cycle slowing at the MBT. We show that the decline of Cdk1 activity releases constraints on Rif1, which then acts to extend the S phase (Summarized in Figure 9). This long S phase is then followed by the first embryonic G2, and a mitosis that is triggered by patterned zygotic transcription of Cdc25. Thus, Rif1 contributes to the developmental program remodeling the embryonic cell cycle.

**Figure 9.**
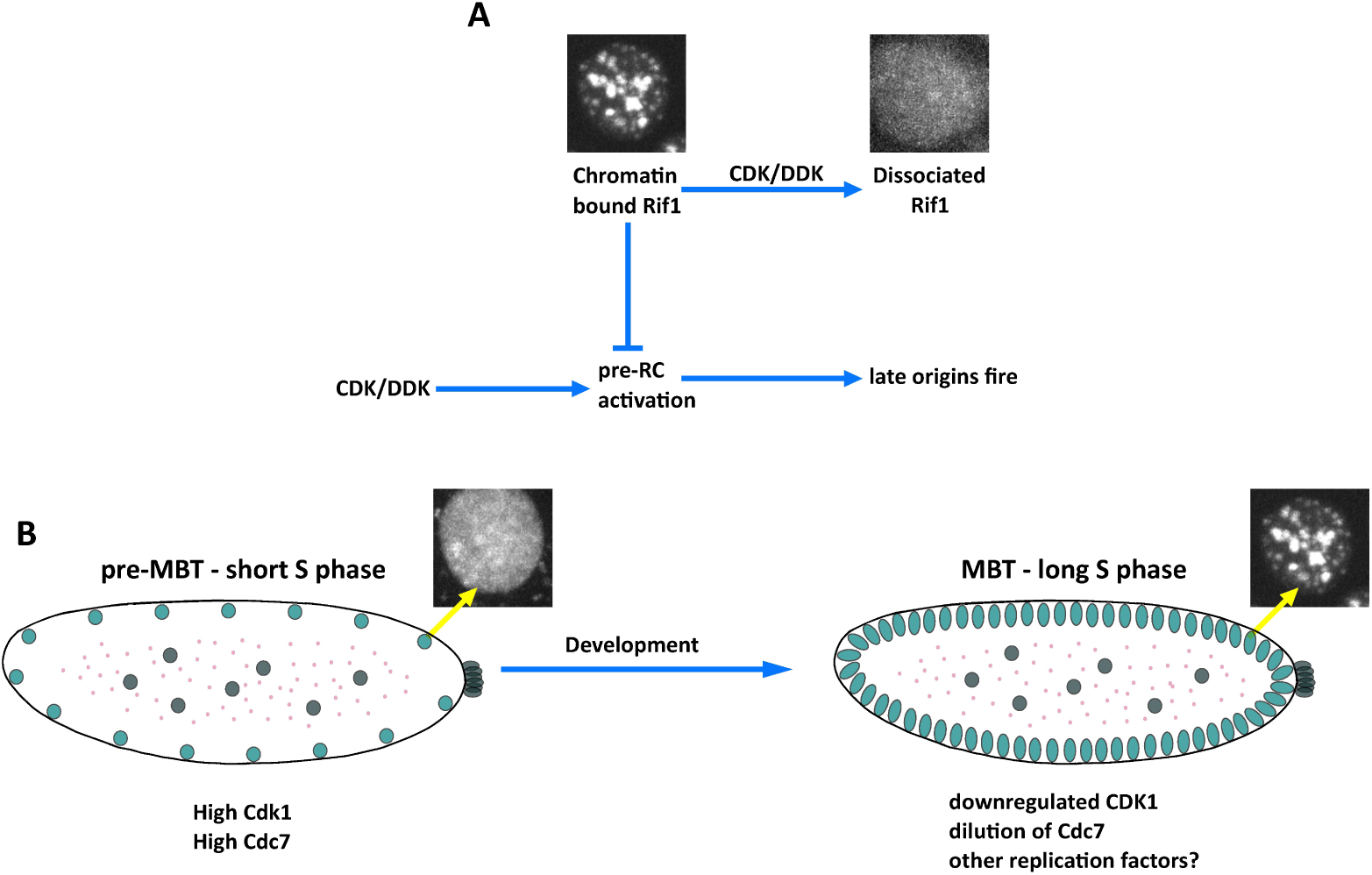
Model. **(A)** Interactions of the S phase kinases and the Rif1 inhibitor create a late replication program. In a nucleus in early S phase (left) foci of compacted late replicating satellites have bound Rif1 that creates a phosphatase rich domain through its interaction with PP1a. This protects the pre-RC from the activating influence of S phase kinases, CDK and DDK. With time (to the right) the S phase kinases act on Rif1, first gradually, to promote its dissociation, and then abruptly as the kinases overwhelm the weakened domain of phosphatase dominance. At this time, the S phase kinases can successfully directly target and activate the pre-RC. **(B)** During the early pre-MBT cycles, high CDK and DDK activities rapidly drive Rif1 dissociation so that delays in replication are initially negligible. A progressive lengthening of the embryonic S phase is associated with incrementally increasing delays in satellite replication. Part of this delay is due to declining CDK and increasing, but still transient, Rif1 association. However, Rif1 independent factors, likely the titration of several replication proteins, such as Cdc7, also contribute to this slowing. Then, at the MBT, developmentally programmed downregulation of Cdk1 allows persistent binding of Rif1 to satellite DNA and the introduction of a late replication program, thereby extending the duration of S phase.

### Isolating a function of Rif1 as a developmental regulator of the replication timing program

Rif1 is widely conserved and it has been implicated in several functions, notably, the control of telomere growth, replication timing, and DNA damage responses (Mattarocci *et al.* 2016). Despite this complexity, the association of Rif1 with replication timing in our studies appears to be uncomplicated by other actions of Rif1. Two factors are likely to contribute to this. First, it is not clear that Rif1 serves the same spectrum of functions in all of the organisms in which it is found. For example, the distinctive telomeres of *Drosophila* do not appear to be regulated by Rif1 (Sreesankar *et al.* 2012). Additionally, previous tests of the function of *Drosophila* Rif1 in cultured cells and when ectopically expressed in other species suggested that *Drosophila* Rif1 protein might lack some functions attributed to the Rif1 of other species (Sreesankar *et al.* 2012). Second, even if the *Drosophila* Rif1 protein has additional functions at other stages in the life of the fly, the first time Rif1 functions in embryogenesis, its role appears to be to impose a replication-timing program on S phase. Thus, this developmental context can isolate the replication-timing role of Rif1 from other potential functions.

### Programmed interaction of Rif1 with satellite sequences guides their timing of replication

#### Developmentally programmed association of Rif1 with satellite sequences marks the onset of late replication

We have previously characterized the program of replication of satellite sequences. Prior to cell cycle 11, when satellite sequences are early replicating, we did not detect significant recruitment of Rif1 to chromatin. In subsequent pre-MBT cycles, when slight and incrementally increasing delays in satellite sequence replication occur, we see brief association of Rif1 to individual blocks of satellite sequence, and an advance in their replication in *rif1* mutant embryos. In pace with the gradual increase in the delay of satellite replication, this Rif1 association persists longer in successive cycles (Figure 1A). In time with the MBT and a dramatic extension of cell cycle 14, we detect more persistent association of Rif1 to satellite sequences.

#### Within each cell cycle, Rif1 dissociation from individual satellite sequences marks the onset of their replication

Individual satellites, each a distinct array of repeats, initiate replication as a unit at a particular time within S phase 14 (Shermoen *et al.,* 2010). The disappearance of different Rif1 foci during S phase parallels the schedule of late replication for the different satellite repeats. Indeed, high-resolution imaging demonstrated that the dissociation of Rif1 is followed closely by the recruitment of PCNA and replication. In addition, TALE-light probes against the satellite 1.686 allowed us to visualize the association of Rif1 with this repeat during S phase. Rif1 staining overlapped that of the satellite early in S phase and Rif1 dissociated in mid S phase just before the decompaction and replication of the underlying DNA (Figure 2). Finally, expression of the Rif1.15a gain-of-function mutant blocked dissociation of Rif1, and prevented complete replication (Figure 5). Thus, the dissociation of Rif1 coincides with and is needed for onset of replication.

#### Phosphorylation controls the association of Rif1 with late replicating chromatin

The control of pre-RC activation by the kinases CDK and DDK appears to be widely conserved among eukarya (Siddiqui *et al.* 2013). The two kinase-types act by parallel mechanisms phosphorylating their respective target sites in the N-terminal regions of MCM2 and MCM4 to promote activation of the pre-RC. Activation usually requires collaboration of the two kinases. These kinases are active from the outset of S phase, suggesting that the timing program of replication within S phase involves local regulation of their action. Rif1 has emerged as a key regulator of this same step and it appears to have two types of interactions with the activating kinases. Rif1 inhibits and delays replication initiation by recruiting PP1 to the genome where its ability to dephosphorylate kinase targets counters the action of DDK and likely CDK. Additionally, DDK and CDK inhibit Rif1, resulting in de-repression that further promotes pre-RC activation (Figure 5 and 7) (Davé *et al.* 2014; Hiraga *et al.* 2014). Opposing actions of Rif1 and DDK were strongly supported by the observation that mutation of *rif1* can suppress the requirement for *cdc7* in yeasts, a genetic interaction that we have now shown is conserved in *Drosophila.*

When Cdk1 activity is especially high at metaphase, Rif1 is absent from chromosomes. After metaphase/anaphase inactivation of Cdk1, Rif1 associates with the separating anaphase chromosomes (Figure 3) (Xu and Blackburn 2004). Manipulations and mutations that promote Cdk1 activity accelerate dissipation of Rif1 from chromatin (Figure 3D, Figure 4). Reciprocally, inhibition of Cdk1 during pre-MBT S-phases increases the amount of time Rif1 spends in foci. Several observations suggest that Cdk1 acts directly on Rif1. Rif1 has conserved clusters of Cdk1 target sites. We observed increased phosphorylation of Rif1 during the early cycles when Cdk1 activity is high and when Rif1 shows minimal association with chromatin. Finally, mutation of 15 CDK and DDK consensus sites in the C-terminus of Rif1 largely prevented dissociation of Rif1 from chromatin. Additionally, DDK also appears to contribute to suppression of Rif1 activity in early embryos. This follows from our finding that Rif1 mutation restored cell cycle progression to embryos blocked by knockdown of DDK (Cdk7). Apparently, Rif1 is active (destructively) in the early embryo without DDK suppression. Thus, both types of S-phase-activating kinase suppress Rif1 activity, a coherent feed-forward input that supports direct action of these kinases to activate the Pre-RC (Figure 9).

In budding and fission yeast, it was suggested that DDK inhibits Rif1 by promoting release of PP1. *Drosophila* Rif1 also interacts with PP1a (Davé *et al.* 2014; Hiraga *et al.* 2014), but we have not detected a change in this interaction when comparing two stages, one in which high Cdk1 prevents Rif1 function and a stage when Cdk1 is inactive (Figure 5). Instead, we see regulation in the association of Rif1 to chromatin. Although we can’t exclude regulation of phosphatase binding, our results demonstrate a key role for phosphorylation-regulated chromatin binding of the *Drosophila* Rif1. Perhaps both mechanisms operate. Since the regulatory architecture is the same, different organisms might emphasize different means of Rif1 inhibition.

The observed program of Rif1 dissociation suggests that the process could play a role in a mysterious feature of replication timing. At least in higher organisms, large domains of the genome replicate as temporal units at a specific time during S phase, a phenomenon that requires coordinate firing of many linked origins (Jackson and Pombo 1998). Concerted dissociation of Rif1 from each satellite sequences is followed by the recruitment of PCNA to the entire domain, suggesting that behavior of Rif1 might coordinate the firing of the numerous origins within each large block of repeating sequence. How might region-specific dissociation of Rif1 itself be coordinated? Our data suggests that dissociation is promoted by phosphorylation of Rif1 (Figure 9). Since Rif1 is known to recruit PP1, the localized phosphatase could counter the action of the kinases, protecting Rif1 from dissociation. This antagonism between kinase activity and localized PP1 could create a bistable circuit. Domains enriched in Rif1 would also be enriched in PP1, which could locally protect Rif1 from the kinases. However, even a slow rate of Rif1 release would eventually give the kinases the upper hand within the domain. At the tipping point dissociation of Rif1 would accelerate. Since our findings show coordinate release of Rif1 from large domains of the genome and show that Rif1 release is coupled to onset of replication, we suggest that the circuit regulating Rif1 release coordinates the firing of origins in local domains.

### S phase extension at the MBT involves interplay of Rif1 and kinases

Our identification of the role of Rif1 in extending S phase and identification of kinase inputs influencing Rif1 activity have implications regarding the control of early embryonic development and mechanisms governing the slowing of the cell cycle at the MBT. In *Drosophila,* the early increases in cell cycle duration are coupled to increases in the time required for S phase (Edgar and O’Farrell 1989, 1990; Sibon *et al.* 1997; McCleland *et al.* 2009; Shermoen *et al.* 2010). This increase in S phase duration is due to the onset of delays in the replication of satellite sequences in S phase (Shermoen et al., 2010). Prior work showed that the activity of Cdk1 influences the delays in satellite replication and the extension of S phase (Farrell et al., 2012; Farrell and O’Farrell, 2013). The speed of the early cycles required high Cdk1, and the abrupt extension of S phase at cycle 14 depended on an MBT associated sharp reduction in interphase Cdk1 activity. The findings reported here show Rif1 associates with satellite sequences and delays their replication upon the downregulation of Cdk1. Thus, the regulatory connection between Cdk1 and Rif1 couples S phase prolongation, and hence cell cycle prolongation with Cdk1 downregulation.

We found that maternally supplied Rif1 is present ubiquitously throughout the pre-MBT cell cycles, and so we suggest that CDK and DDK kinases are critical to prevent maternally supplied Rif1 from interfering with rapid S phase completion in early cycles. There is progressive reduction of Cdk1 activity in successive interphases prior to the MBT (Edgar et al., 1994) that allows limited Rif1 association to satellites during cycles 11-13, and this association persists longer in each cycle. Cdk1 activity is normally required during these S phases for their timely completion, and we show that this requirement can be bypassed by mutation of *rif1* (Figure 6). However, while the abrupt cycle-14 extension of S phase required *rif1,* only a portion of the incremental and progressive extension of S phase required *rif1* and gradual prolongation of S phase still occurred in *rif1* mutant embryos, suggesting that there are other inputs.

### Rif1 independent contributions to S phase duration

In *rif1* embryos, the early 3.4 min S phase is extended to 27 min by cycle 14. Although much shorter than the 72 minute S phase 14 in wild-type embryos, this rif1-independet extension of S phase resembled progression normally seen in the early cycles. S phase extended progressively in small but increasing increments and replication of satellite sequences was selectively delayed. These observations show that the distinctive behavior of satellite sequences is not entirely dependent on Rif1 interaction, and that additional factors contribute to S phase extension during the progressive phase.

The progressive delays in timing of replication of satellite sequences in *rif1* mutant *Drosophila* embryos suggest that selective delays in the activation of replication are involved. A long-standing suggestion for embryonic slowing of the cell cycle is that the increasing number of nuclei titrate factors required for the speedy early S phases. Several factors might limit pre-RC activation. For example, Cdc7, the kinase component of activating DDK, has an input as demonstrated by its genetic interaction with Rif1. Given that we saw diminishing concentration of nuclear Cdc7 as it was distributed to an increasing large number of nuclei over successive divisions, it is possible that developmental declines in this activating input could contribute to slowing of S phase. However, beyond the factors we have discussed — Rif1, Cdk and DDK — there are other inputs. Notably, the concentrations of several proteins that are recruited to the pre-RC during its maturation influence the speed and efficiency of pre-RC activation in yeast and in *Xenopus* (Mantiero *et al.* 2011; Collart *et al.* 2013).

Investigations of S phase extension occurring at the *Xenopus* MBT have emphasized multiple inputs influencing activation of the pre-RC (Walter 1997; Collart *et al.* 2013). A recent analysis suggested that Drf1, the activating subunit for the Cdc7 kinase in the *Xenopus* embryo, is subject to active destruction at the MBT (Collart *et al.* 2017). Failure to down regulate DDK sensitized embryos to the further stress of overproduction of three replication factors that promote pre-RC maturation. Indeed, the authors suggest that titration of the maternal supply of these three replication proteins cooperate with active destruction of Drf1 to slow S phase at the MBT. Thus, evolution appears to have brought the same regulatory step under developmental control in very different organisms, but many factors complicate a comparison between systems, and here we emphasize that the detail of the dissection of the process in *Drosophila* has led us to a specific conclusion regarding Rif1 control of cell cycle slowing at the MBT in this organism.

In contrast to the early progressive slowing of the cell cycle in *Drosophila* embryos, the abrupt extension of cell cycle 14 shows features of a distinct regulatory transition. It has switch-like features (Lu *et al.* 2009). The triggering of this sudden extension of S phase is not simply due to a limitation of a factor, because, unlike the earlier progressive extension of S phase, inhibition of transcription prevents the abrupt extension (Shermoen *et al.* 2010). This shows that existing materials are adequate to support another relatively fast cycle and that the extension of cycle 14 involves an active process. And importantly, we have previously shown that the key event in triggering the extension is the downregulation of cyclin:Cdk1 (Farrell *et al.* 2012; Farrell and O’Farrell 2013). Here, we show that it also depends on the positive action of Rif1 as a repressor of replication and that this activity of Rif1 depends on the downregulation of Cdk1. In summary we suggest that there are two distinct phases of S phase prolongation operating in the *Drosophila* embryo, one progressive and earlier and one abrupt and occurring at the onset of cycle 14. The *rif1* mutant is defective in the abrupt event that marks the MBT.

### Implications of the dispensability of Rif1

We found it surprising that *rif1* mutants have a highly penetrant phenotype with a specific failure to prolong S phase 14, but yet produce viable progeny. This means that the prolongation of S phase 14 is not essential and that subsequent contributions of *rif1* to development and survival are also dispensable. We would like to put this in context.

Rif1 loss is not without major consequences. Both hatching of maternally deficient embryos and survival of zygotically deficient flies are compromised. So Rif1 is important, but a detailed characterization will be required to understand its contributions to survival. Nonetheless, it is clear that mutant embryos with a 27 min S phase 14 instead of the normal S phase duration of more than an hour can hatch. *rif1* mutant embryos still undergo an MBT and the majority of the embryos slow their cell cycle because they arrest in a G2 after their abnormally fast S phase 14. A minority of the embryos undergoes a complete or partial extra syncytial cycle suggesting that S phase timing contributes to regulation that normally introduces a G2 into the cell cycle at the MBT with great reliability.

The late replication program in cell cycle 14 precedes obvious appearance of heterochromatic marks on satellite sequences (Yuan and O’Farrell 2016). These marks do appear later in cell cycle 14 and their appearance was not compromised in the absence of Rif1. Thus, neither the selective localization of Rif1 to satellite sequences nor the specific time of replication of the satellites in cycle 14 are required to trigger or guide the appearance of the heterochromatic marks. Importantly, we showed previously that recruitment of HP1a and associated acquisition of other heterochromatin marks delays replication of the X chromosomal 359-satellite repeat in cycle 15 (Yuan et al., 2016). Here we show that in the absence of Rif1, the 359 repeat still exhibits late replication in cycle 15, suggesting that heterochromatin can act by a Rif1 independent pathway to cause replication delay. While *rif1* mutant embryos may show subtle alterations in the program of late replication in later cycles, presently the unique dependency of replication timing on Rif1 function appears limited to a narrow window of developmental time. Thus, survival of *rif1* mutants does not assess the importance of late replication per se, only the impact of the delay in cycle 14.

### Concluding remarks

Embryonic development presents early cell cycles with an unusual challenge – regulating cell division in the absence of transcription. Perhaps because of this early constraint, the biology of early embryos is streamlined and lacks many regulatory processes that appear later in development. When development introduces complications, it does so incrementally, highlighting individual regulatory circuits. By focusing on early development, we have been able to isolate and study the contributions of Rif1 to late replication and to S phase length. Our work uncovers a special developmental function for Rif1 in controlling the timely prolongation of S phase at the Mid Blastula Transition, and provides new insight into the control of late replication.

## MATERIALS AND METHODS

### Fly stocks

All *Drosophila melanogaster* stocks were cultured on standard cornmeal-yeast medium. The following fly strains were used in this study: w^1118^ Canton-S (wild type), his2av-mRFP (Bloomington stock number 23650 or 23651), mRFP-HP1a (30562), eyeless-gal4 (5535), maternal tubulin-gal4 (7063), UAS-shRNA^cdc7^ (TRiP GL00585), mei41^D3^ (gift from Tin Tin Su, University of Colorado Boulder), mCherry-pcna (this study), rif1-EGFP (this study), cdc7-EGFP (this study), rif1^-^ (this study), UASp>Rif1-3xFlag (this study), UASp>Rif1-15a-3xFlag (this study).

### Generation of transgenic lines described in this study

All transgensis injections were performed by Rainbow Transgenic Flies, Camarillo, CA.

The stocks rif1-EGFP and cdc7-EGFP were generated using CRISPR-Cas9 editing to modify the endogenous rif1 and cdc7 genes (diagrammed in Supplemental Figure 1) as described in Gratz et al 2014. Briefly, a single CRISPR target site was selected as close to the stop codon as possible. To direct homologous repair, approximately 1.5 kb of DNA on either side of the cut site was amplified from gDNA isolated from the vasa-Cas9 stock (51324). Both homology arms, a 6xGlyGlySer-EGFP tag, and the endogenous 3’UTR were cloned into the pHD-DsRed vector using Gibson Assembly. To generate guide RNA plasmids, annealed DNA oligos were cloned into the pU6-BbsI-chiRNA vector. The donor plasmid and the gRNA plasmid were coinjected into vasa-cas9 embryos by Rainbow Transgenic Flies. After screening for DsRed^+^ progeny, successful modification was confirmed by PCR genotyping and anti-GFP immunoblotting. A single male containing a successful modification was backcrossed to our lab’s w1118 Canton-S stock for five generations to establish a stock.

CRISPR-Cas9 editing was used to generate the rif1 null allele (diagrammed in Supplemental figure 6) by replacing the rif1 ORF with a visible 3xP3-DsRed marker. Two CRISPR target sites were selected, one site directly upstream of the start codon and one site 14 bp upstream of the stop codon. Gibson assembly was used to construct a homologous repair donor plasmid containing approximately 1.5 kb of DNA homologous to the genomic sequence directly upstream of the first cut site, the DsRed marker, and approximately 1.5 kb of DNA homologous to the genomic sequence directly downstream of the second cut site. Both gRNA plasmids and the donor plasmid were co-injected into vasa-cas9 (51324) embryos by Rainbow Transgenic Flies. After screening for DsRed^+^ progeny, successful replacement of the rif1 ORF was confirmed by PCR genotyping (Supplemental figure 6) and by Sanger sequencing across the recombinant breakpoints. A single male carrying the rif1 null allele was backcrossed to our lab’s wild-type strain for five generations to establish the rif1^-^ stock.

To generate the mCherry-pcna transgenic stock approximately 1 kb of sequence upstream of the pcna start codon, the mCherry-5xGlyGlySer tag, and the pcna gene were cloned into the pw+attB vector using Gibson Assembly. The resulting transgenesis plasmid and phiC31 mRNA were co-injected into attP-9a embryos (9744) to perform phiC31 mediated integration.

To generate the UAS-Rif1.WT-3xFlag and UAS-Rif1.15a-3xFlag transgenic lines either the rif1 ORF was amplified from embryo cDNA and into the pENTR-D vector by TOPO cloning. In order to generate the phosphomutant Rif1.15a, a DNA fragment containing the mutated sequence was synthesized (Integrated DNA Technologies, Redwood City, CA) and used to replace the corresponding wild-type sequence in the ENTRY plasmid by Gibson Assembly. The ENTRY vectors were then recombined with the pPWF-attB construct using the LR Clonase II kit (Invitrogen, Carlsbad, CA). The resulting transgenesis plasmids and phiC31 mRNA were co-injected into attP-9a embryos (9744) to perform phiC31 mediated integration.

Plasmids and DNA sequences are available upon request.

### Embryo staining

Embryos were collected on grape agar plates, washed into mesh baskets and dechorinated in 50% bleach for 2 minutes. Embryos were then devitellenized and fixed in a 1:1 mixture of methanol-heptane, before storing in methanol at −20°C. Embryos were gradually rehydrated in a series of increasing PTx:Methanol mixtures (1:3, 1:1, 3:1) before washing for 5 minutes in PTx (PBS with 0.1% Triton). Embryos were then blocked in PBTA (PTx supplemented with 1% BSA and 0.2% Azide) for 1 hour at room temperature. Blocked embryos were then incubated with the primary antibody overnight at 4°C. The following primary antibodies were used: rabbit anti-GFP at 1:500 (Invitrogen A11122), and mouse anti-tubulin at 1:100 (DSHB 12G10, AA12.1-s, and AA4.3-s). Embryos were then washed with PTx for three times 15 minutes each, and then incubated with the appropriate fluorescently labeled secondary antibody (Molecular Probes) at 1:300 for 1 hour in the dark at room temperature. Embryos were then washed again with PTx for three times 15 minutes each. In order to detect total DNA, DAPI was added to the second wash. Finally stained embryos were mounted on glass slides in Fluoromount.

To detect both Rif1-EGFP and the AATAC satellite repeat, formaldehyde fixed embryos were blocked in PBTA for 1 hour, incubated with rabbit anti-GFP at 1:500 (Invitrogen A11122) for 1 hour at room temperature, washed three times for 15 minutes each in PTX, incubated with fluorescently labeled secondary antibody at 1:300 for 1 hour at room temperature in the dark, and then washed three times for 15 minutes in PTX. Embryos were then post-fixed in 4% formaldehyde for 20 minutes. FISH was then performed as described in Shermoen et al., 2010 using a Cy5 labeled AATAC 30-mer oligo probe (IDT).

### Protein extract preparation and immunoprecipitation

Protein extracts were prepared by homogenizing embryos in embryo lysis buffer (50 mM Tris-HCl pH 8.0, 100 mM NaCl, 1% Triton X-100, 1 mM EDTA) supplemented with 1X protease inhibitor cocktail (Pierce), PMSF, 100 mM Sodium Fluoride, 100 mM β-gylcerophosphate, and 10 mM sodium pyrophosphate. After lysis, 2X Sample Buffer was added to the extract, and samples were boiled for 5-10 minutes. For dephosphorylation reactions protein extracts were prepared as above except for the omission of phosphatase inhibitors. Extracts were then supplemented with 1 mM MnCl_2_ and incubated with ƛ phosphatase for 30 minutes at 30°C. The reaction was terminated by adding 2X Sample Buffer and boiling for 5-10 minutes.

For immunoprecipitation of GFP tagged Rif1 dechorinated embryos were lysed using a dounce homogenizer in 500 μl of ice cold RIPA buffer (50 mM Tris-HCl pH 8.0, 150 mM NaCl, 1% Triton X-100, 0.5% Sodium deoxycholate, 0.1% SDS, 1 mM DTT) supplemented with 1X protease inhibitor cocktail (Pierce, Waltham, MA), PMSF, 100 mM Sodium Fluoride, 100 mM β-gylcerophosphate, and 10 mM sodium pyrophosphate. The resulting extract was cleared by spinning twice at 12,000 rpm for 10 minutes each at 4°C, after which the supernatant was incubated with 20 μl of GFP-TRAP Magnetic beads (Chromotek, Martinsried, Germany) for 1 hour at 4°C on a nutator. The beads were then washed for three times with lysis buffer using a magnetic rack, and proteins were eluted by boiling in 30 μl of 2x Sample Buffer.

### Western Blotting

For standard western blotting, protein extracts were separated by electrophoresis in precast 4-15% polyacrylamide gels (Biorad, Hercules, CA). To separate phosphorylated forms of Rif1, protein extracts were run on a 0.5% agarose strengthened 3% polyacrylamide gel containing 20 μM Mn^2+^-Phostag (AAL-107, Wako Chemicals, Japan). Proteins were then transferred to a PVDF membrane using a wet transfer system. The membrane was blocked in TBST (TBS with 0.1% Tween-20) supplemented with 2.5% BSA and then incubated in the appropriate primary antibody at a dilution of 1:1000 overnight at 4°C. The blot was then washed three times for 10 minutes each in TBST, and then incubated in the appropriate secondary antibody at a dilution of 1:10,000 for 1 hour at room temperature. The blot was then washed three times for 10 minutes each in TBST, and then treated with Pierce SuperSignal West Pico ECL, and used to expose autoradiography film. The following antibodies were used: rabbit anti-GFP (ab290, Abcam, Cambridge, United Kingdom), rabbit anti-PP1a (2582, Cell Signaling, Danver, MA), Goat anti-Rabbit HRP conjugated (Biorad).

### Microinjection

Embryo microinjections were performed as described in Farrell et al., 2012. In vitro transcribed mRNA was prepared using the CellScript T7 mRNA production system (CellScript, Madison, WI) as described in Farrell., et al 2012 and injected at a concentration of 600 ng/μl. The purified proteins used in this study were described in Yuan and O’Farrell, 2016.

### Microscopy

For live imaging embryos were collected on grape agar plates, and aged at 25°C when appropriate. After dechorination in 50% bleach, embryos were aligned and glued to glass coverslips, and then covered in halocarbon oil before imaging. Embryos were imaged using a spinning disk confocal microscope, and the data were analyzed using Volocity 6 (Perkin Elmer, San Jose, CA). For most experiments, approximately 30 embryos were watched under the microscope, and only the embryos at the appropriate developmental stage were filmed. When comparing different conditions, all images acquired in a single experiment were acquired at the same time and with identical microscope settings. Choice of objective and z-stack size were determined by experimental need. Unless otherwise noted all images shown are projections of a z-stack series. When imaging embryos following microinjection, the objective was centered in the portion of the embryo nearest the site of injection to record the area of maximum effect.

### Reproducibility

The development of embryos is highly stereotyped and the quality of the data is founded in the resolution of the imaging. Still images and movies presented in this paper are representative of multiple embryos filmed during a single experiment (technical replicates) and of identical experiments performed on embryos collected from independently sorted flies (biological replicates). Choice of embryos for imaging was dictated by developmental stage and by embryo health (unfertilized and clearly damaged embryos were excluded). Western blots reported in this paper were performed in two biological replicates (separate protein extracts from independent embryo collections).

## Acknowledgements

We thank past members of the O’Farrell laboratory, especially Kai Yuan, Antony Shermoen and Mark McCleland, for guidance and reagents. We also thank Danielle O’Farrell for early work on this project. The Bloomington stock center and *Drosophila* community provided important reagents throughout this work. This research was supported by National Institutes of Health grant GM037193/R37 (to P.H.O), NIH Grant GM007810/T32, and an ARCS Foundation Fellowship (to C.A.S).

## Supplemental Figures

**Supplemental Figure 1.**
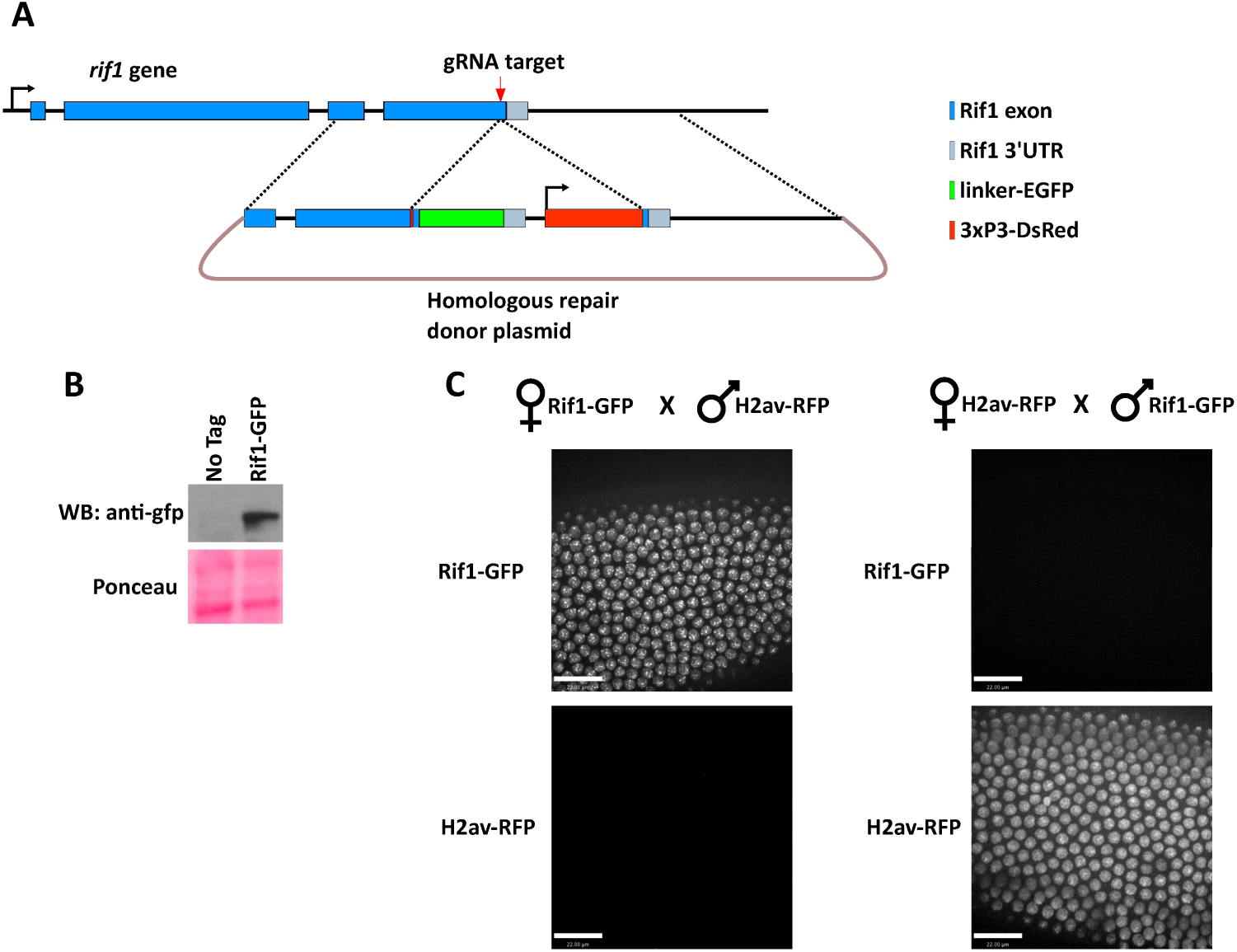
Generation and characterization of endogenously tagged Rif1::EGFP flies. **(A)** Schematic showing the gene structure of *drif1* and the CRISPR-Cas9 tagging strategy. Briefly, a single guide RNA was chosen to direct Cas9 cleavage in the extreme C-terminus of the rif1 ORF. A donor plasmid containing approximately 1.5 kb of homology to either side of the break point was used to insert a GlyGlySer (linker)-EGFP tag in addition to a DsRed selectable maker under the control of the eye specific 3xP3 enhancer. **(B)** Anti-GFP western blot on embryonic protein extract to confirm successful tagging. **(C)** Still frames from time-lapse confocal imaging of embryos produced from the indicated crosses. Selected images are from individual embryos at 10 minutes into interphase of cycle 14. The Rif1 protein present at the MBT is maternally provided.

**Supplemental Figure 2.**
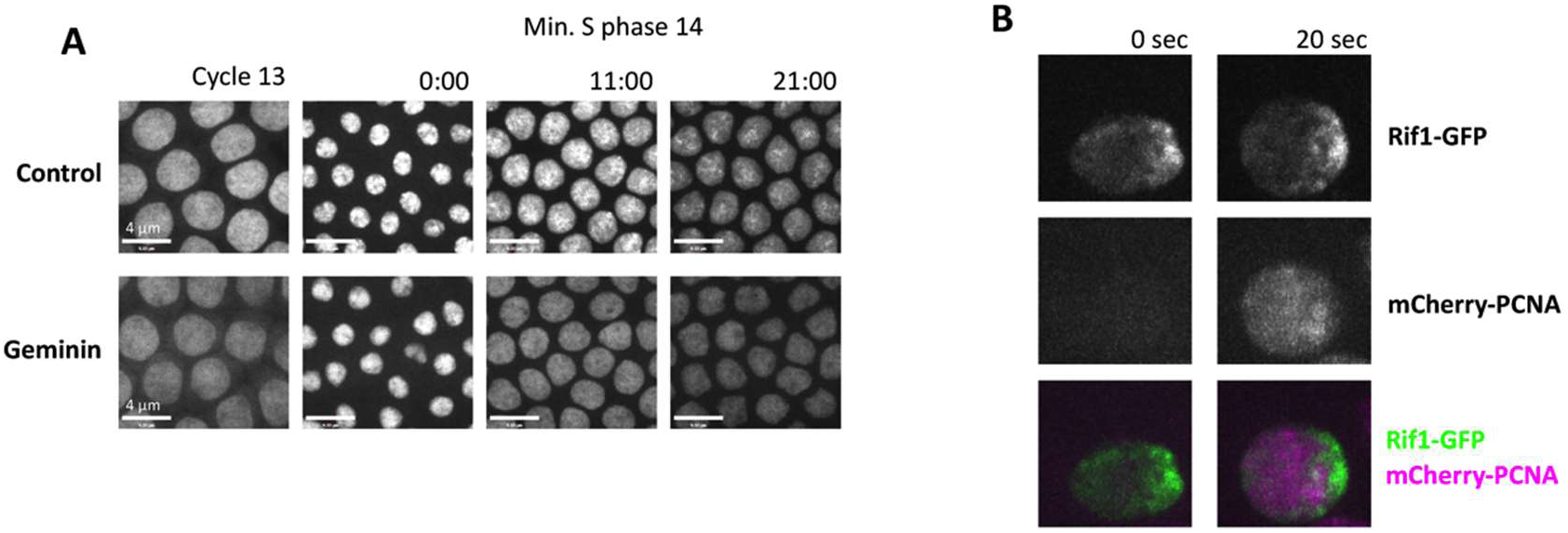
**(A)** Injection of geminin eliminates S phase 14 foci of mCherry-PCNA. In control embryos, mCherry-PCNA marks nuclear locations of active DNA replication resulting in bright PCNA foci later in S phase. When pre-RC formation is blocked by the injection of purified geminin protein during interphase 13 the nuclear PCNA signal is overall less intense, and never resolves into replication foci. We conclude that transgenic mCherry-PCNA faithfully marks replicating sequences. **(B)** Stills from time-lapse imaging of Rif1-EGFP and mCherry-PCNA during the start of S phase 15. Note that the recruitment of Rif1 precedes the recruitment of PCNA to chromatin. Once S phase begins PCNA is spread throughout the early replicating euchromatic portion of the nucleus, but the PCNA signal does not overlap with the Rif1 bound late replicating heterochromatin which by cycle 15 is concentrated to one edge of the nucleus.

**Supplemental Figure 5.**
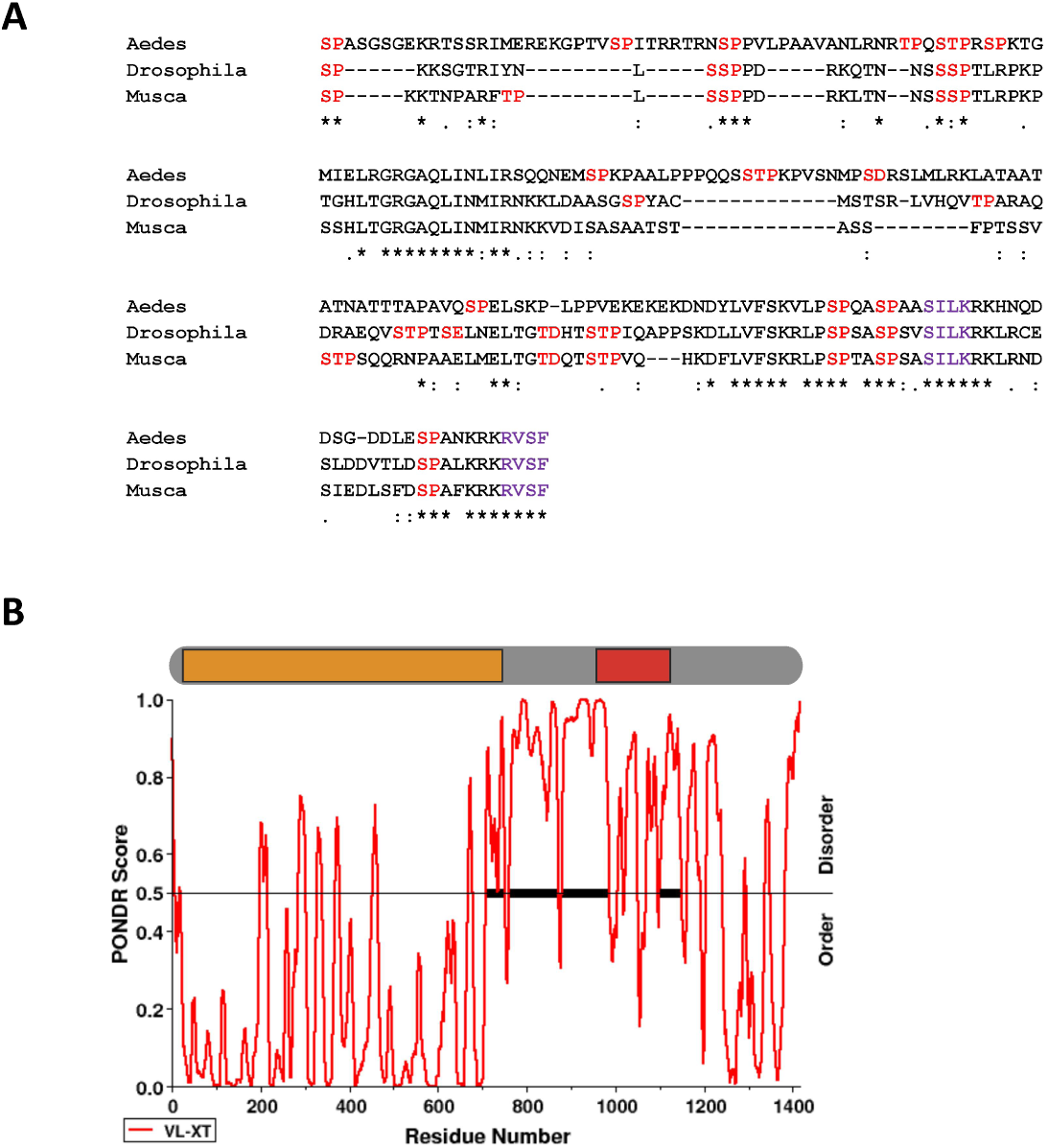
Analysis of potential Rif1 CDK and DDK phosphorylation sites. **(A)** Multiple sequence alignment of the indicated portions of the Rif1 protein sequences from *Aedes aegypti* (1345-1541), *Drosophila melanogaster* (946-1103), and *Musca domestica* (1041-1189) using Clustal Omega. Potential DDK and CDK phosphorylation sites are highlighted in red. Both CDK and DDK are serine/threonine kinases where specificity is encoded by the residue in the +1 position. CDK phosphorylates S/T residues followed by a proline. DDK targets S/T residues followed by an acidic group which can be provided by an acidic amino acid (D or E) or by a previous phosphorylation (e.g. SSP). PP1 interaction motifs are highlighted in purple. **(B)** Analysis of the *D. melanogaster* Rif1 protein sequence using the Predictor of Natural Disordered Regions (PONDR) tool to score for regions of intrinsic disorder. PONDR scores above 0.5 suggest regions of intrinsic disorder. Above the graph is a schematic of the relevant regions of the Rif1 protein. The N-terminal heat repeats are represented by the yellow box, and the portion Rif1 containing the potential CDK and DDK phosphorylation sites analyzed in **(A)** is represented by the red box.

**Supplemental Figure 6.**
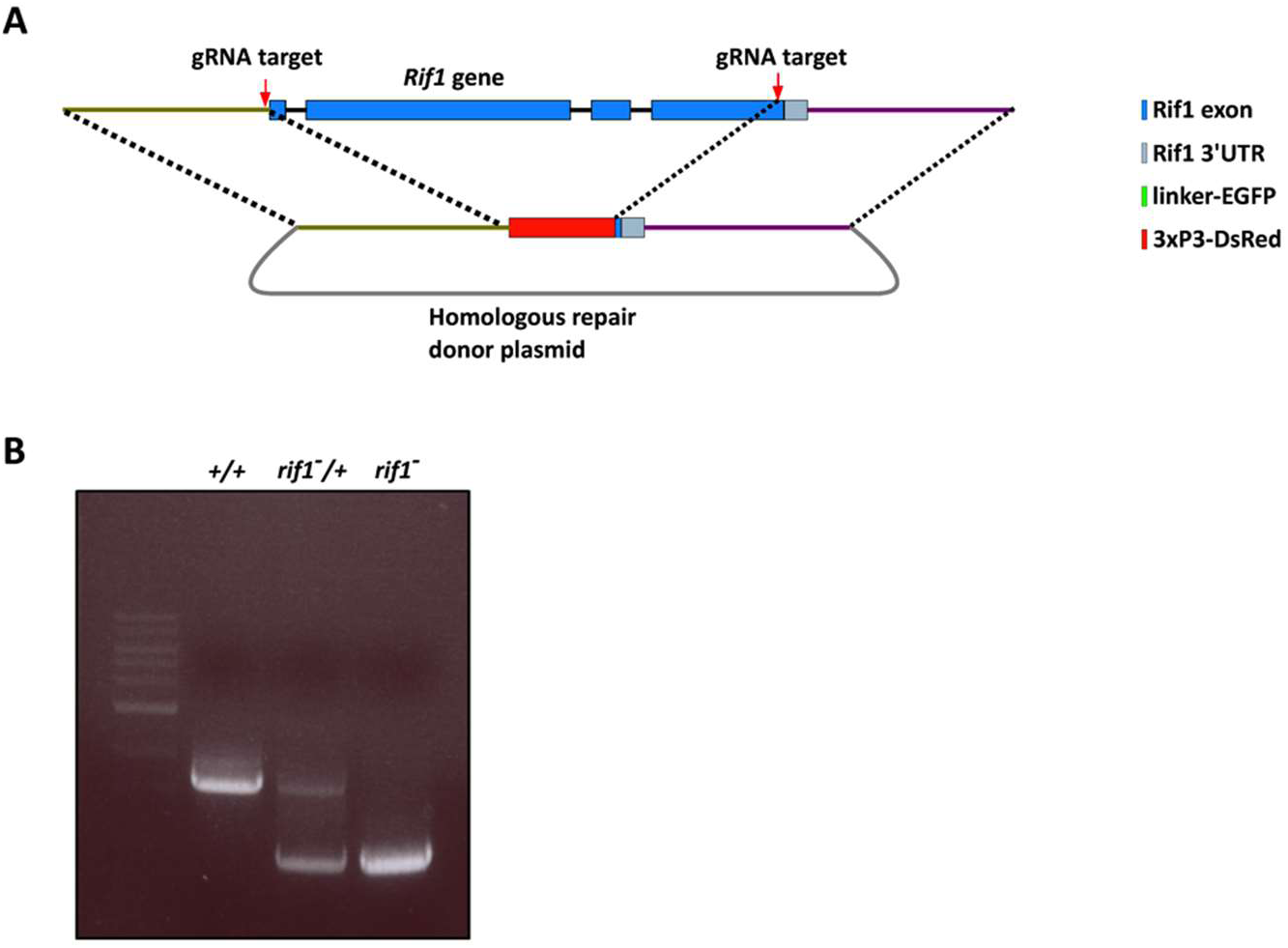
(A) Schematic showing the gene structure of drif1 and the CRISPR-Cas9 editing strategy used to generate the *rif1* null allele. Briefly, two CRISPR target sites were selected, one site directly upstream of the start codon, and one site directly upstream of the stop codon. Approximately 1.5 kb of DNA homologous to the genomic sequence on either upstream or downstream of the break points was used to direct the replacement of the rif1 ORF with the visible 3xP3-DsRed marker. (B) Confirmation of correct replacement of the rif1 ORF by PCR.

## Movie legends

**Movie 1 Rif1 forms dynamic nuclear foci as the embryonic cell cycle lengthens**. This video accompanies Figure 1D. Live imaging of a Rif1-GFP (green) and H2aV-RFP (magenta) in an embryo completing the blastoderm cell cycles and the MBT. Video begins at mitosis 10 and ends following mitosis 14 in domain 5. Note that foci of Rif1 grow in number and duration as the cell cycle slows, and that Rif1 foci disappear progressively as interphase completes. Z stacks were acquired every 2 minutes on a 100x oil objective.

**Movie 2 Imaging the disappearance of Rif1 foci**. This video accompanies Figure 1E. Live imaging of Rif1-GFP at high frame rate (every 20 seconds) during S phase 13. When Rif1 dissociates from the chromatin the focus of Rif1-GFP erodes from the outside over the course of 2 minutes as the signal fades.

**Movie 3 Coordination of Rif1 dissociation and DNA replication**. This video showing Rif1-GFP (green) and mCherry-PCNA (purple) during S phase 14 accompanies Figure 2A. Movie starts at the end of S phase 13 and follows nuclei through S phase 14. Rif1 dissociates from chromatin before the underlying sequences acquire PCNA signal indicating onset of late replication. Absence of coincidence of the two signals is evident throughout the sequence. While complicated by nuclear movements, recruitment of PCNA to previously Rif1 staining regions is apparent near the close of S phase. Z stacks were acquired every 2 minutes on a 100x oil objective.

**Movie 4 Rif1 is recruited to the late replicating heterochromatin during Mitosis 14 before the beginning of S phase 15**. This video accompanies Figure 2B. Live imaging of Rif1-GFP (green) and mCherry-PCNA (magenta) during mitosis 14 and the start of S phase 15. Rif1 binds to the chromatin during mitotic exit, and is enriched at the leading edge of the chromosome mass where the late replicating pericentric heterochromatin resides. Rif1 binding is evident before the appearance of PCNA signal on the chromatin, which marks the start of S phase. Once S phase begins, the Rif1 stained heterochromatin does not overlap with the PCNA stained early replicating euchromatin. Z stacks were acquired every 20 seconds on a 100x oil objective.

**Movie 5 Imaging Rif1 and the replication of the 1.686 satellite repeat**. This video accompanies Figure 2C. Live imaging of Rif1-GFP (green) and the late replicating satellite 1.686 using an mCherry labeled TALE-light protein (magenta) during S phase 13. Before 1.686 replicates, it is marked by Rif1 (coincident signal appears white). Rif1 then dissociates (at 00:04:20) before this repeat decondenses (at 00:04:40) and replicates. After replication, the TALE-light signal appears brighter and recompacted (00:09:00), but lacks a Rif1 signal. Finally, the 1.686 loci separate, align, and then lose fluorescence upon progression to mitosis. Z stacks were acquired every 20 seconds on a 100x oil objective.

**Movie 6 Ectopic activation of Cdk1 during S phase 14 drives Rif1 dissociation and accelerates late replication**. This video accompanies Figure 4B. Live imaging of Rif1-GFP (green) and mCherry-PCNA (magenta) during S phase 14 following injection of Cdc25 mRNA during cycle 13. Introduction of this Cdk1 activator accelerates the loss of Rif1 foci and drives the underlying sequences to replicate earlier in S phase (compare to movie 3). Z stacks were acquired every 2 minutes on a 100x oil objective.

**Movie 7 *rif1* embryos undergo an early mitosis 14**. This video accompanies Figure 6C. Live imaging of GFP-HP1a injected into in a wild-type embryo (left) and a *rif1* embryo (right). Movies begin in cycle 12 and follow embryos through cycle 14. Note that the *rif1* embryo undergoes an extra synchronous mitosis approximately 23 minutes into interphase 14. Z stacks were acquired at minute and twenty second intervals on a 20x air objective.

**Movie 8 Level of nuclear Cdc7 decline over the blastoderm cycles leading up to the MBT**. This video accompanies Figure 7B. Live imaging of Cdc7-GFP (green) and His2av-RFP (magenta) during cell cycles 11 through 14. Z stacks were acquired every minute on a 20x air objective.

## REFERENCES

Aggarwal B. D., Calvi B. R., 2004 Chromatin regulates origin activity in Drosophila follicle cells. Nature 430: 372–376.

Alver R. C., Chadha G. S., Gillespie P. J., Blow J. J., 2017 Reversal of DDK-Mediated MCM Phosphorylation by Rif1-PP1 Regulates Replication Initiation and Replisome Stability Independently of ATR/Chk1. Cell Rep 18: 2508–2520.

Baldinger T., Gossen M., 2008 Binding of Drosophila ORC proteins to anaphase chromosomes requires cessation of mitotic cyclin-dependent kinase activity. Molecular and Cellular … 29: 140–149.

Blumenthal A. B., Kriegstein H. J., Hogness D. S., 1974 The units of DNA replication in Drosophila melanogaster chromosomes. Cold Spring Harb Symp Quant Biol 38: 205–223.

Collart C., Allen G. E., Bradshaw C. R., Smith J. C., Zegerman P., 2013 Titration of four replication factors is essential for the Xenopus laevis midblastula transition. Science 341: 893–896.

Collart C., Smith J. C., Zegerman P., 2017 Chk1 Inhibition of the Replication Factor Drf1 Guarantees Cell-Cycle Elongation at the Xenopus laevis Mid-blastula Transition. Dev Cell 42: 82–96.

Cornacchia D., Dileep V., Quivy J.-P., Foti R., Tili F., Santarella-Mellwig R., Antony C., Almouzni G., Gilbert D. M., Buonomo S. B. C., 2012 Mouse Rif1 is a key regulator of the replication-timing programme in mammalian cells. EMBO J 31: 3678–3690.

Davé A., Cooley C., Garg M., Bianchi A., 2014 Protein phosphatase 1 recruitment by Rif1 regulates DNA replication origin firing by counteracting DDK activity. Cell Rep 7: 53–61.

Deneke V. E., Melbinger A., Vergassola M., Di Talia S., 2016 Waves of Cdk1 Activity in S Phase Synchronize the Cell Cycle in Drosophila Embryos. Dev Cell 38: 399–412.

Dimitrova D. S., Gilbert D. M., 2000 The spatial position and replication timing of chromosomal domains are both established in early G1 phase. Mol Cell 4: 983–993.

Edgar B. A., O’Farrell P.H., 1989 Genetic control of cell division patterns in the Drosophila embryo. Cell 57: 177–187.

Edgar B.A., O’Farrell P.H., 1990 The three postblastoderm cell cycles of Drosophila embryogenesis are regulated in G2 by string. Cell 62: 469–480.

Farrell J. A., O’Farrell P. H., 2013 Mechanism and regulation of Cdc25/Twine protein destruction in embryonic cell-cycle remodeling. Curr Biol 23: 118–126.

Farrell J. A., O’Farrell P. H., 2014 From egg to gastrula: how the cell cycle is remodeled during the Drosophila mid-blastula transition. Annu Rev Genet 48: 269–294.

Farrell J. A., Shermoen A. W., Yuan K., O’Farrell P. H., 2012 Embryonic onset of late replication requires Cdc25 down-regulation. Genes Dev 26: 714–725.

Graham, C. F. and Morgan, R. W., 1966 Changes in the cell cycle during early amphibian development. Dev Biol 14: 439–460.

Hayano M., Kanoh Y., Matsumoto S., Renard-Guillet C., Shirahige K., Masai H., 2012 Rif1 is a global regulator of timing of replication origin firing in fission yeast. Genes Dev 26: 137–150.

Hiraga S.-I., Alvino G. M., Chang F., Lian H.-Y., Sridhar A., Kubota T., Brewer B. J., Weinreich M., Raghuraman M. K., Donaldson A. D., 2014 Rif1 controls DNA replication by directing Protein Phosphatase 1 to reverse Cdc7-mediated phosphorylation of the MCM complex. Genes Dev 28: 372–383.

Hiratani I., Ryba T., Itoh M., Yokochi T., Schwaiger M., Chang C.-W., Lyou Y., Townes T. M., Schübeler D., Gilbert D. M., 2008 Global reorganization of replication domains during embryonic stem cell differentiation. PLoS Biol 6: e245.

Holt L. J., Tuch B. B., Villén J., Johnson A. D., Gygi S. P., Morgan D. O., 2009 Global analysis of Cdk1 substrate phosphorylation sites provides insights into evolution. Science 325: 1682–1686.

Jackson D. A., Pombo A., 1998 Replicon clusters are stable units of chromosome structure: evidence that nuclear organization contributes to the efficient activation and propagation of S phase in human cells. J Cell Biol 140: 1285–1295.

Knott S. R. V., Peace J. M., Ostrow A. Z., Gan Y., Rex A. E., Viggiani C. J., Tavaré S., Aparicio O. M., 2012 Forkhead transcription factors establish origin timing and long-range clustering in S. cerevisiae. Cell 148: 99–111.

Kumar S., Yoo H. Y., Kumagai A., Shevchenko A., Shevchenko A., Dunphy W. G., 2012 Role for Rif1 in the checkpoint response to damaged DNA in Xenopus egg extracts. Cell Cycle 11: 1183–1194.

Lima-De-Faria A., 1959 Differential uptake of tritiated thymidine into hetero- and euchromatin in Melanoplus and Secale. J Cell Biol 6: 457–466.

Lima-De-Faria A. and Jaworska H., 1968 Late DNA synthesis in heterochromatin. Nature 217: 138–142.

Labib K., 2010 How do Cdc7 and cyclin-dependent kinases trigger the initiation of chromosome replication in eukaryotic cells? Genes Dev 24: 1208–1219.

Lu X., Li J. M., Elemento O., Tavazoie S., Wieschaus E. F., 2009 Coupling of zygotic transcription to mitotic control at the Drosophila mid-blastula transition. Development 136: 2101–2110.

Makunin I. V., Volkova E. I., Belyaeva E. S., Nabirochkina E. N., Pirrotta V., Zhimulev I. F., 2002 The Drosophila suppressor of underreplication protein binds to late-replicating regions of polytene chromosomes. Nature 160: 1023–1034.

Mantiero D., Mackenzie A., Donaldson A., Zegerman P., 2011 Limiting replication initiation factors execute the temporal programme of origin firing in budding yeast. EMBO J 30: 4805–4814.

Mattarocci S., Hafner L., Lezaja A., Shyian M., Shore D., 2016 Rif1: A Conserved Regulator of DNA Replication and Repair Hijacked by Telomeres in Yeasts. Front Genet 7: 45.

Mattarocci S., Shyian M., Lemmens L., Damay P., Altintas D. M., Shi T., Bartholomew C. R., Thomä N. H., Hardy C. F. J., Shore D., 2014 Rif1 controls DNA replication timing in yeast through the PP1 phosphatase Glc7. Cell Rep 7: 62–69.

McCleland M. L., Shermoen A. W., O’Farrell P. H., 2009 DNA replication times the cell cycle and contributes to the mid-blastula transition in Drosophila embryos. J Cell Biol 187: 7–14.

Milán M., Campuzano S., Garcia-Bellido A. 1996 Cell cycling and patterned cell proliferation in the Drosophila wing during metamorphosis. Proc Natl Acad Sci USA 93: 640–645.

Neufeld T. P., la Cruz de A. F., Johnston L. A., Edgar B. A., 1998 Coordination of growth and cell division in the Drosophila wing. Cell 93: 1183–1193.

Raff J. W., Glover D. M., 1988 Nuclear and cytoplasmic mitotic cycles continue in Drosophila embryos in which DNA synthesis is inhibited with aphidicolin. J Cell Biol 107: 2009.

Raghuraman M. K., Brewer B. J., Fangman W. L., 1997 Cell cycle-dependent establishment of a late replication program. Science 276: 806–809.

Rhind N., Gilbert D. M., 2013 DNA replication timing. Cold Spring Harb Perspect Biol 5: a010132.

Schübeler D., Scalzo D., Kooperberg C., van Steensel B., Delrow J., Groudine M., 2002 Genome-wide DNA replication profile for Drosophila melanogaster: a link between transcription and replication timing. Nat Genet 32: 438–442.

Shermoen A. W., McCleland M. L., O’Farrell P. H., 2010 Developmental control of late replication and S phase length. Curr Biol 20: 2067–2077.

Sheu Y.-J., Stillman B., 2010 The Dbf4-Cdc7 kinase promotes S phase by alleviating an inhibitory activity in Mcm4. Nature 463: 113–117.

Sibon O. C., Stevenson V. A., Theurkauf W. E., 1997 DNA-replication checkpoint control at the Drosophila midblastula transition. Nature 388: 93–97.

Siddiqui K., On K. F., Diffley J. F. X., 2013 Regulating DNA replication in eukarya. Cold Spring Harb Perspect Biol 5: a012930.

Snow, MHL. 1976. “Embryo growth during the immediate postimplantation period.” CIBA Foundation Symposium 40. 53–70.

Sreesankar E., Bharathi V., Mishra R. K., Mishra K., 2015 Drosophila Rif1 is an essential gene and controls late developmental events by direct interaction with PP1-87B. Sci Rep 5: 10679.

Sreesankar E., Senthilkumar R., Bharathi V., Mishra R. K., Mishra K., 2012 Functional diversification of yeast telomere associated protein, Rif1, in higher eukaryotes. BMC Genomics 13: 255.

Stephenson R., Hosler M. R., Gavande N. S., Ghosh A. K., Weake V. M., 2014 Characterization of a Drosophila ortholog of the Cdc7 kinase: a role for Cdc7 in endoreplication independent of Chiffon. J Biol Chem 290: 1332–1347.

Stepinska U. and Olszanska B., 1983 Cell multiplication and blastoderm development in relation to egg envelope formation during uterine development of quail *(Coturnix coturnix japonica)* embryo. Dev Biol 228: 505–510.

Su T. T., O’Farrell P. H., 1997 Chromosome association of minichromosome maintenance proteins in Drosophila mitotic cycles. J Cell Biol 139: 13–21.

Tanaka S., Araki H., 2013 Helicase activation and establishment of replication forks at chromosomal origins of replication. Cold Spring Harb Perspect Biol 5: a010371.

Tanaka S., Nakato R., Katou Y., Shirahige K., Araki H., 2011 Origin association of Sld3, Sld7, and Cdc45 proteins is a key step for determination of origin-firing timing. Curr Biol 21: 2055–2063.

Tanaka S., Umemori T., Hirai K., Muramatsu S., Kamimura Y., Araki H., 2006 CDK-dependent phosphorylation of Sld2 and Sld3 initiates DNA replication in budding yeast. Nature 445: 328–332.

Taylor J. H., 1960 Nucleic acid synthesis in relation to the cell division cycle. Ann N Y Acad Sci 90: 409–421.

Vogelauer M., Rubbi L., Lucas I., Brewer B. J., Grunstein M., 2002 Histone acetylation regulates the time of replication origin firing. Mol Cell 10: 1223–1233.

Walter J., 1997 Regulation of Replicon Size in Xenopus Egg Extracts. Science 275: 993.

Xu L., Blackburn E. H., 2004 Human Rif1 protein binds aberrant telomeres and aligns along anaphase midzone microtubules. J Cell Biol 167: 819–830.

Yamazaki S., Ishii A., Kanoh Y., Oda M., Nishito Y., Masai H., 2012 Rif1 regulates the replication timing domains on the human genome. EMBO J 31: 3667–3677.

Yeeles J. T. P., Deegan T. D., Janska A., Early A., Diffley J. F. X., 2015 Regulated eukaryotic DNA replication origin firing with purified proteins. Nature 519: 431–435.

Yuan K., O’Farrell P. H., 2016 TALE-light imaging reveals maternally guided, H3K9me2/3-independent emergence of functional heterochromatin in Drosophila embryos. Genes Dev 30: 579–593.

Yuan K., Seller C. A., Shermoen A. W., O’Farrell P. H., 2016 Timing the Drosophila Mid-Blastula Transition: A Cell Cycle-Centered View. Trends Genet 32: 496–507.

Yuan K., Shermoen A. W., O’Farrell P. H., 2014 Illuminating DNA replication during Drosophila development using TALE-lights. Curr Biol 24: R144–5.

Zegerman P., Diffley J. F. X., 2006 Phosphorylation of Sld2 and Sld3 by cyclin-dependent kinases promotes DNA replication in budding yeast. Nature 445: 281–285.

Zimmermann M., Lottersberger F., Buonomo S. B., Sfeir A., de Lange T., 2013 53BP1 regulates DSB repair using Rif1 to control 5’ end resection. Science 339: 700–704.

Zofall M., Smith D. R., Mizuguchi T., Dhakshnamoorthy J., Grewal S. I. S., 2016 Taz1-Shelterin Promotes Facultative Heterochromatin Assembly at Chromosome-Internal Sites Containing Late Replication Origins. Mol Cell 62: 862–874.

